# Community review: a robust and scalable selection system for resource allocation within open science and innovation communities

**DOI:** 10.1101/2022.04.25.489391

**Authors:** Chris L. B. Graham, Thomas E. Landrain, Amber Vjestica, Camille Masselot, Elliot Lawton, Leo Blondel, Luca Haenel, Bastian Greshake Tzovoras, Marc Santolini

## Abstract

Resource allocation is essential to the selection and implementation of innovative projects in science and technology. With large stakes involved in concentrating large fundings over a few promising projects, current “winner-take-all” models for grant applications are time-intensive endeavours that mobilise significant researcher time in writing extensive project proposals, and rely on the availability of a few time-saturated volunteer experts. Such processes usually carry over several months, resulting in high effective costs compared to expected benefits. Faced with the need for a rapid response to the Covid19 pandemic in 2020, we devised an agile “community review” system to allocate micro-grants for the fast prototyping of innovative solutions. Here we describe and evaluate the implementation of this community review across 147 projects from the “Just One Giant Lab’s OpenCOVID19 initiative” and “Helpful Engineering” open research communities. The community review process uses granular review forms and requires the participation of grant applicants in the review process. Within a year, we organised 7 rounds of review, resulting in 614 reviews from 201 reviewers, and the attribution of 48 micro-grants of up to 4,000 euros. We show that this system is fast, with a median process duration of 10 days, scalable, with a median of 4 reviewers per project independent of the total number of projects, and fair, with project rankings highly preserved after the synthetic removal of reviewers. We investigate the potential bias introduced by involving applicants in the process, and find that review scores from both applicants and non-applicants have a similar correlation of r=0.28 with other reviews within a project, matching previous observations using traditional approaches. Finally, we find that the ability of projects to apply to several rounds allows to both foster the further implementation of successful early prototypes, as well as provide a pathway to constructively improve an initially failing proposal in an agile manner. Overall, this study quantitatively highlights the benefits of a frugal, community review system acting as a due diligence for rapid and agile resource allocation in open research and innovation programs, with particular implications for decentralised communities.

## Introduction

The distribution of scientific funding through grants requires the identification of novel, feasible and potentially impactful projects. However, the traditional scientific grant allocation system involving a closed panel of experts in the field, or in similar fields (1), is notoriously slow (2), time consuming and expensive, often taking months and occurring in timescales of yearly rounds or grant calls. In extreme cases, the grant review program can be more costly than simply allocating small grants to each applicant, as in the case of the NSERC grant system of 2008 (3). In addition, the allocation of grants has shown to suffer from various biases, such as the composition of the grant panel (4), gender and geographical location (5), group based dynamics personality triumphing over other qualitative factors (6–8), and socio-psychological factors such as group dynamics and personality traits triumphing over other qualitative factors (8,9). Overall, selection results are only weakly predictive of future performance (10).

Often, the reason to conduct grant allocations in a ‘closed’ setting is to protect the intellectual property of the grant applicants. As a result, the majority of unsuccessful grant applications, which contain a large amount of research effort, are inevitably lost, unavailable to the public after the fact (11). The recent emergence of the open science movement (12–14) has reversed this incentive, with open access practices and early sharing of results such as pre-registration now becoming normalised by institutions and journals (15).

Beyond the allocation of funding, the review of early-stage, unpublished work by community peers has been leveraged to allocate other types of resources. For example, conferences often need to allocate time for their participants to showcase their work to other members of the community during a usually short amount of time, thereby providing a platform for promoting the work, building novel collaborations, and getting feedback to improve a manuscript. In such cases, peer reviewing is needed to decide in a collegial fashion whether a work is worth a full oral presentation, a shorter lightning talk, a poster, or is not of a high enough standard to be showcased to participants. For example, the EasyChair online platform has been used by close to 100k conferences for handling such review processes (16). Often, participants to a conference are also part of the “program committee” reviewing the proposed abstracts and papers of peer applicants, alongside external members of the scientific community. This allows for a rapid process usually lasting less than a few weeks.

This suggests there is a potential for a new, more agile route for community-driven grant allocation bypassing pre-selected grant panels that handle funds and introduce barriers (8), and relying instead on peer applicants to handle a large-scale application process in a short timescale. In this study, we present the design, implementation, and results of a community-driven, open peer-review system to support two open research communities during the COVID pandemic across seven selection rounds (**Fig1**): the “OpenCOVID19” initiative from Just One Giant Lab (JOGL) (14,17) and the COVID relief charity Helpful Engineering (18). We show that this system is robust (unaffected by reviewer removal), agile (fast timeline), iterative (covering multiple grant rounds), decentralised (driven by the community), and scalable. Finally, we discuss these results and the perspectives they offer for the design of future community-driven review systems.

**Figure 1.**
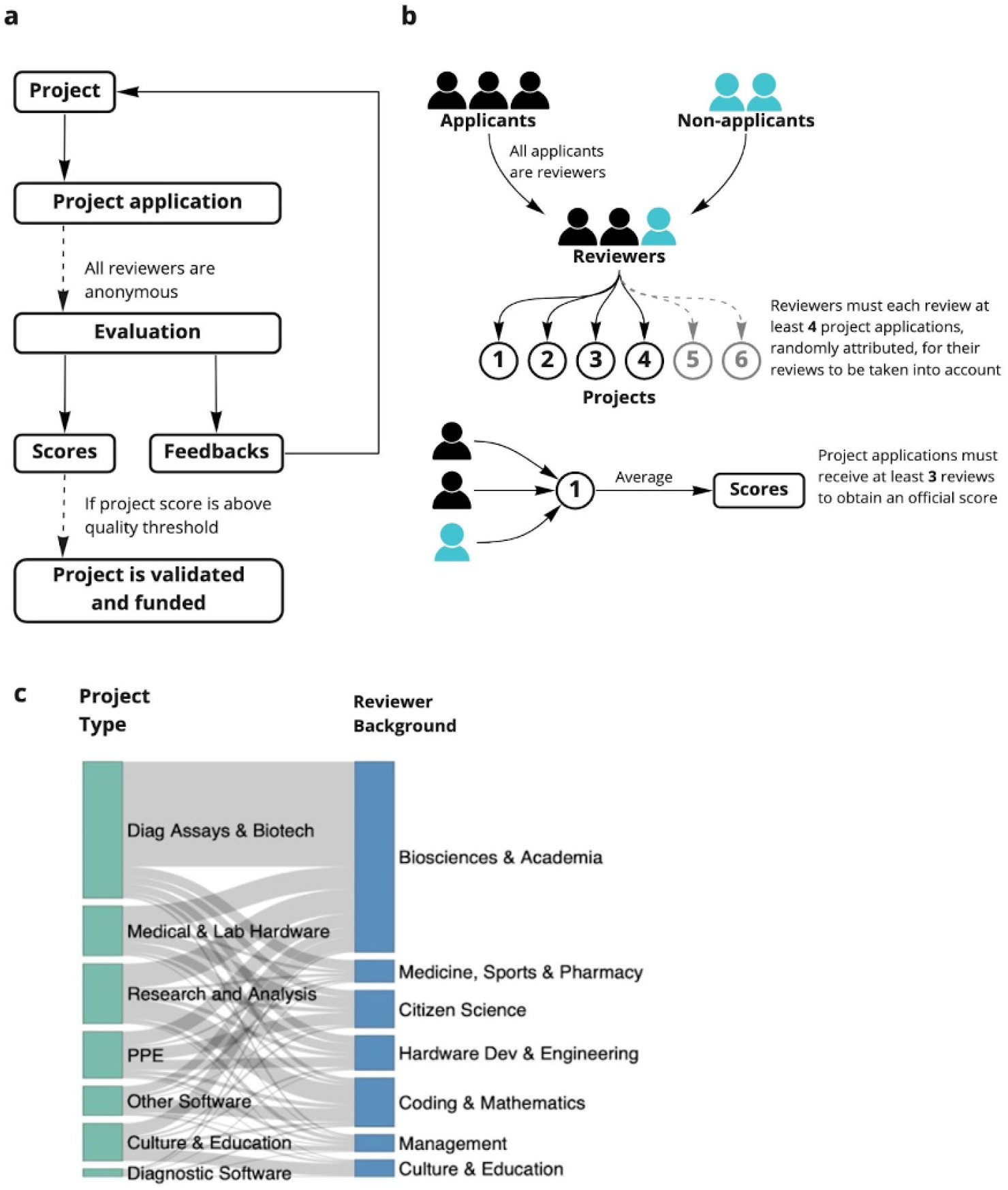
Overview of the open peer review process. **a** Stages of the open peer review process JOGL rounds 3-5. The online review forms and templates are found in supplementary data. **b** community review method JOGL rounds 3-5 **c** distribution of project type to expertise across rounds

## Methods

### Context

The implementation of a crowd-based, open peer-review system followed the need to support two nascent community efforts, first by allocating volunteers to projects in the COVID relief charity Helpful Engineering (18), then by allocating funding to projects in the JOGL “OpenCOVID19” initiative (17). The method was developed as an open access grant review and funding allocation system, meaning that it was open to anyone willing to review. It was implemented using the Just One Giant Lab platform (app.jogl.io) as the project proposal host, and free-to-use tools and forms to conduct the review process (**FigS2**). The implementation was applied and refined over 7 rounds across 1 year.

### General process of review

The peer review system was conducted on early phase projects within both JOGL and Helpful Engineering. These projects were submitted by project leaders to a grant review process in order to allocate volunteers in the case of Helpful Engineering, and funding in the context of OpenCovid19. Reviews of these projects (see **Fig1b**) were initially conducted by members of the community and included members of other projects who also submitted their project for review.

As a consequence of the process being experimental and serving an urgent need, the process was altered over time. However it followed the same general pattern (**Fig1, FigS1**). First, a template for the grant proposal was created by the community and was iteratively edited (Supplementary material). This template followed typical grant application templates (19), with sections on team composition, the project general hypothesis and its timeline. The proposal was then submitted using a google form, which requested an email address and required only one application per project (**FigS2a**). In Helpful Engineering rounds this included a link to their proposal hosted in editable google documents, while in JOGL rounds this included instead a link to their open access JOGL page proposal. The project links were manually formatted into a google sheet with a link to a review form for convenience, along with descriptions of desirable reviewer skills by the applicants in the proposal submission form to help reviewers find relevant projects (**FigS2B**). A technical evaluation form scoring various criteria (eg: proposal efficacy, team composition, impact) on a scale from 1-5 (**Supplementary Information**) was created by the designers of the program and iteratively changed following feedback from the community (**FigS2c**). This form separated questions on projects into two areas centred around Impact and Feasibility for ease of identifying the problems and/or strengths in their grant application. A message with a link to the reviewer form for use in review, along with a nested google sheet containing project proposal links was spread among the community through announcements and email. In later rounds (JOGL 3-5) all applicants were asked to review at least three other projects and the process was randomised, removing the need for a sheet. The review process was given between 4 days HE 1, 8 days HE 2, 7 Days - JOGL 1, 10 days - JOGL 2, 16 days - JOGL 3, 21 days - JOGL 4 and 28 days JOGL 5, (**FigS1b**) to allow reviews to occur and be collected via a google form into a google sheet automatically (**FigS2d**). No reviewer selection was performed, however usernames (Slack handles or JOGL user names depending on the round) and emails were collected for conducting further analyses. The average reviewer scores were then composed into a presentation to the community, and those projects with a score above a given impact/feasibility threshold (**FigS2e**) were chosen for grant funding. Due to the community aspect of our study, members from the JOGL HQ participated in the process, and their removal from the analysis does not change the observations (**FigS10**), we therefore retain these in our analysis.

### Iterative changes to the review process

As mentioned in the previous section, the method of review was iteratively changed throughout the programme, elongating from an initial “emergency style” four day period of review and allocation (HE round 1) to 21 and 28 days in JOGL rounds 4 and 5 as the need for rapid response reduced, with an overall median average of 10 days per round (**FigS1b**). As such, the design of the general process described in **Fig1** and **FigS1** had some variations. For example, initially applicants were not required to review applications (**Fig1b**). Upon scaling up of the programme, the process was adapted to be less dependent on volunteer reviewers, (**Fig S1b,A-D**) and more dependent on the applicant’s reviews of their competing peers (**Fig1c**). In JOGL rounds 3, 4 and 5 (**FigS1b**) teams depositing a proposal could only be eligible after having reviewed at least three other teams. The changes in the process and differences in the rounds are summarised in **FigS1c**. The major changes between Helpful Engineering (HE) and JOGL rounds (**FigS1c**) occurred through changes in the nature of proposal submission from google document links to an online project repository. In addition, HE rounds offered no grants, but instead publicity and allocation of members to projects, while JOGL offered microgrants worth up to 4000 euros per team (**FigS2c**).

### Final selection process

In Helpful Engineering, this review method allowed 54 projects to be reviewed and ranked by score for community recruitment purposes, with no official threshold, but instead an arbitrary set of “Highlighted projects”. Within JOGL this grant system reviewed 96 eligible applications (**Fig2**) and allocated requested funds to 36 of these. Once the review process had taken place, the cut-off threshold of scores given by reviewers to projects for funding by JOGL was decided by an absolute threshold (above 3.5/5 average reviewed score) rather than a rejection rate. The absolute 3.5/5 threshold was chosen due to the gap in project scores in the first JOGL round, and maintained at this standard for consistency. Those with a score above the threshold were funded.

**Figure 2.**
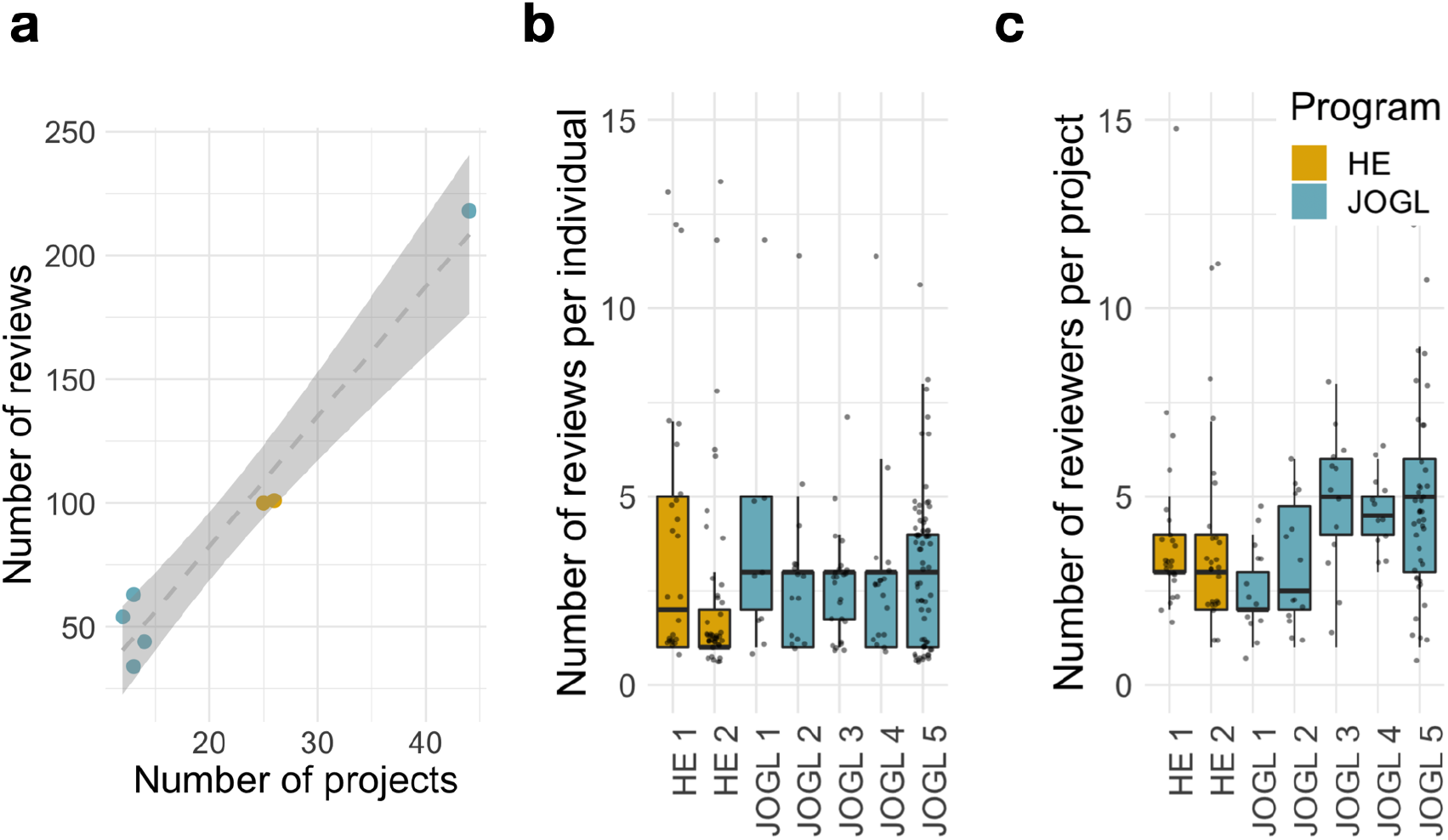
Scalability of the community review methodology. **a** Number of Reviewers and projects during each round of peer/grant review. HE-Helpful Engineering Crowd reviews, JOGL-Just One Giant Lab funded projects. **b** Number of reviews per individual reviewer. **c** Number of reviewers per project. Despite a scale-up in the number of projects, the number of reviews per round scales linearly with the number of projects applying.

### Detection of fraudulent reviewer behaviour

The results of each round, and number of reviews per reviewer were closely monitored through simple email handle tracking by a data handling administrator. If a number of emails were found to be grading a particular project and not others this was suggestive of fraudulent behaviour and self-grading. These reviews were then removed, and teams that were found responsible for this bad behaviour were removed from the review process, as described in grant round participation rules. This was performed only one time across all rounds prior to the rule of each reviewer having a minimum review count for their scores to be counted, which was created after this event.

### Computation of inter-review correlations

In order to compute the correlation between reviews within a project, we first proceeded with data cleaning. Indeed, in several rounds, reviewers had to answer only a subset of questions from the review form that corresponded to the topic of the project (e.g data project vs bio project). However, in some cases, projects were assigned to one or the other category by the different reviewers, leading them to answer to different sets of questions, making the correlation only partial. To mitigate this effect, for each project we kept only the reviews that corresponded to the choice of topic that was most expressed among reviewers. If no majority could be found, the project was removed from analysis. We then converted review scores into vectors of length the number of grades in the form. A Spearman’s rho correlation was then computed between all pairs of reviews within a project. Finally, for each review we computed the average correlation with the other reviews in the project. This number was then associated with the features of the reviewer who produced the review (**Fig4 and FigS7**).

### Reviewer feasibility and impact scores

For JOGL rounds 1-5, we categorised the 23 to 29 questions from the review forms into either impact or feasibility related questions (see **Supplementary Data Review forms)**. The feasibility and impact categories were used to provide two dimensional projections of project scores during the result presentation.

### Reviewer professions and project types

For all JOGL rounds, reviewer responses of the “What is your expertise relevant to this project” question were manually coded into simple categories per review (see **Table S1**). This data was then used as a proxy for expertise distribution across rounds **(Figure 1b)**.

In addition, reviewer responses to the “Which category would you say the project falls under?” question were manually coded into a set of simple categories, representing a summary of the project types across rounds per review (see **Supplementary information** conversion table). The data, due to suggested categories provided by the form, needed little manual coding, but was formatted into a list, then concatenated into similar project types for simplicity. This data was used to assess project type distribution across rounds **(Figure 1b)**.

### Bootstrap analysis

In order to perform the bootstrap analysis of **Fig3d**, we first ranked all projects using their average review score across reviewers. We then selected a review at random. If the corresponding project had at least another review, we removed the selected review and recomputed the average scores and final ranking. We then computed the Spearman correlation between the obtained scores and the original scores. This process was repeated until each project had only one review. Finally, we reiterated this analysis 50 times.

**Figure 3.**
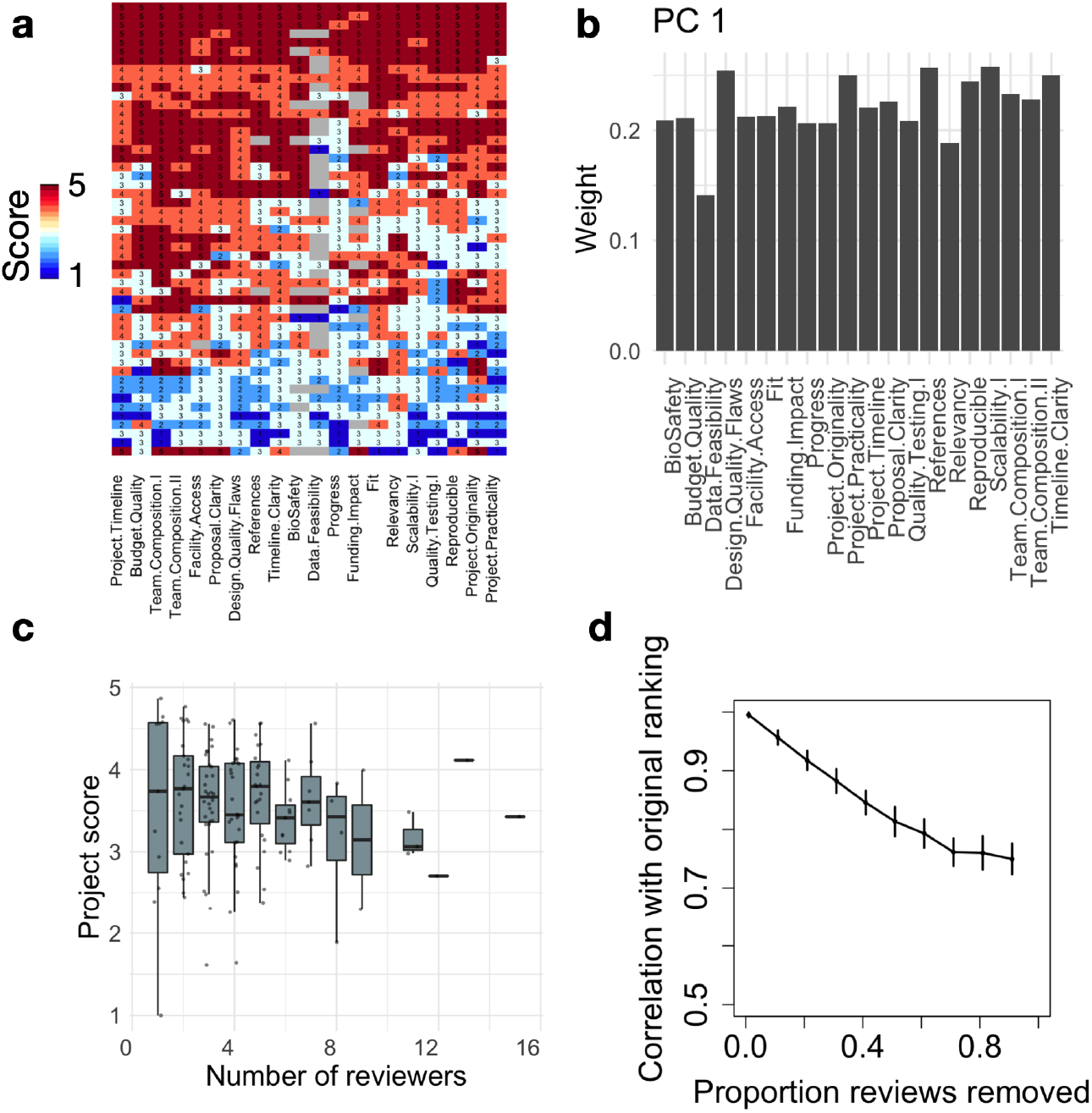
Robustness of the Community review process. **a** Heatmap showing review scores (rows) across questions (columns) for the JOGL round 4. Row and column clustering was performed using correlation distance and average linkage. **b** We show for PC1 (53% variance) the weights of the questions from the original question space. PC1 has near uniform weights across dimensions, indicating that it corresponds to an average score. **c** Project average score across reviewers as a function of number of reviewers. **d** Bootstrap analysis showing the Spearman correlation between the final project ranking and simulated project rankings with increasing proportion of reviews removed from the analysis (see Methods).

## Results

### Scalability of the review process

We describe in **Fig2** the reviewing activity across the seven rounds implemented. Despite the large differences in number of projects between rounds, we find that the number of reviews per round scales linearly with the number of projects applying (**Fig2a**). In addition, the number of reviews per individual and number of reviewers per project have relatively stable distributions across rounds, independent of scale (**Fig2b-a**). For example, despite the substantial growth in reviewers and projects in JOGL round 5, we find that the distributions of number of reviews per reviewer and number of reviewers per project are comparable to those observed in the previous rounds, highlighting the scalability of this review system to different systems. Finally, we note that the number of reviewers per project show a sustained increase from JOGL round 3 onwards, corresponding to the change in review process, where applicants were required to review at least 3 other projects (see Methods). This highlights the benefits of this requirement in promoting sustained engagement.

### Robustness of the final project ranking

In order to obtain a granular score for each project, the reviewers had to grade between 23 (JOGL 1-2) and 29 (JOGL 3-5) criteria in the review form. We first investigate whether these questions would cover different dimensions of project quality. We show in **Fig3a** a heatmap of reviewer scores in JOGL round 4 across 20 questions (removing questions only representing a minority of projects), visually showing a greater inter-review variability (rows) than inter-questions variability (columns). As such, respondents seem to assign a project with either low scores or high scores throughout their review. To quantify the number of dimensions of variation across grades, we conduct a Principal Component Analysis (PCA) on the questions correlation matrix, i.e correlations between pairs of questions across reviews (see **Fig S2a**). We find that the first principal component (PC1) explains most of the variance (53%), with the next largest PC explaining less than 6% of the variance (**Fig S3**). When examining the weights of the various questions in PC1, we find that they all contribute to a similar level (**Fig3b**), meaning that the PC1 is close to the average over all questions, confirming the visual insight from **Fig3a**. This shows that scores are highly correlated, and that the average score across the review form is a reasonable operationalisation of project quality. In addition, we find that the top 10 PCs explain ∼90% of the variance, indicating that review forms could be reduced in complexity using only half of the number of questions to obtain a similar outcome.

We next investigate the reliability of the review scores obtained across reviewers. As suggested by the previous section, for each review we compute the average score across all criteria from the review form. In the following, we refer to this average score as the review score. We observe a generally good discrimination of review scores between projects, with intra-project variation smaller than inter-project variation (**FigS4)**.

Finally, we investigate the robustness of the final project ranking as a function of the number of reviews performed using a bootstrap analysis (see Methods). For each project, a project score is computed by averaging its review scores, and projects are then ranked by decreasing score. We show in **Fig3d** the Spearman correlation between the original project ranking and the ranking obtained when removing a certain proportion of reviews. We find that even with only one review per project, the final ranking is strongly conserved (rho=0.75 and **see FigS5**), confirming that intra-project variability is much smaller than the range of inter-project variability. This supports our design strategy, showing that the use of a granular form allows us to differentiate between projects whilst minimising the impact of individual reviewers variability.

### Measuring reviewer biases

The previous results show the existence of variability between reviews from different reviewers, yet with limited impact on final rankings (**Fig3d**). Here we investigate the source of review variability: is it due to inherent grading variability between individuals, or can it be attributed to other factors? To evaluate this question, we analyse how review score varies with reviewer attributes. We explore in particular two possible sources of bias for which we could gather data: expertise and application status. First, reviewer expertise might be important in determining an accurate project score. This feature is operationalised using the self-reported expertise grade (1 to 5) present in the review forms of JOGL rounds. Second, a majority of reviewers (65%) were applicants of other competing projects, which could lead to a negative bias when reviewing other competing projects.

We show in **Fig4** how the review score varies as a function of these reviewer characteristics. We find that review score increases slightly with expertise (**Fig4a**, Spearman’s rho=0.1, p=0.039). However, the strongest effect is found when looking at applicant bias: review scores from applicants are significantly lower than those from non-applicants (**Fig4b**, p=1.4e-7). Given the fact that in JOGL rounds 3-5, applicants were required to score at least 3 projects, they are found to have a lower expertise towards other projects (**Fig S6**), which could explain the lower scores as suggested by **Fig4a**. Yet, when controlling for review expertise, we find that application status is the main contributing factor, with a score difference between applicants and non-applicants of −0.52 points (p=1.61e-6, **Supplementary Table 1**). This supports that application status is a significant source of bias in the final score.

**Figure 4.**
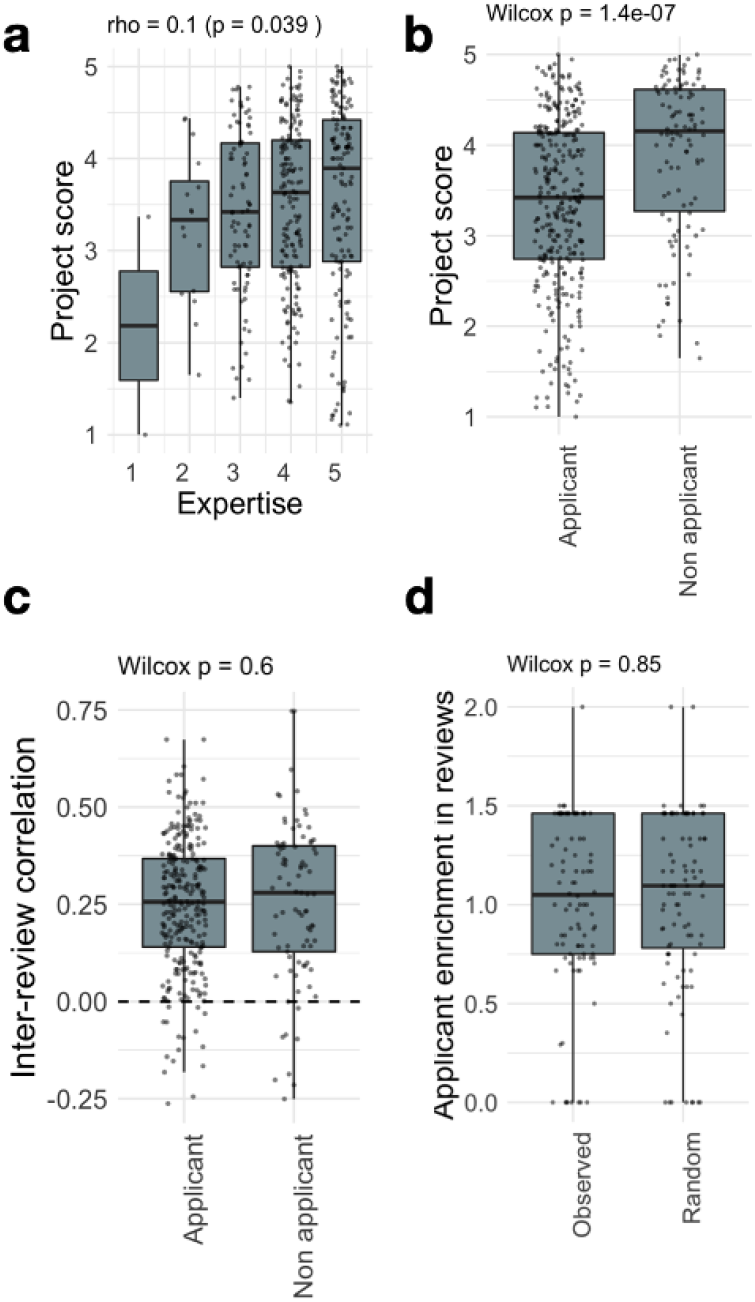
Questionnaire granularity allows to measure and mitigate reviewer biases. Breakdown of project score as a function of **a**. self-assessed expertise, **b**. applicant status (i.e. the reviewer is also an applicant in the round). See Fig S4 for a breakdown by review round. **c**. For each project, we compute the ratio between the proportion of applicant reviewers to the average proportion of applicant reviewers observed in the round. The boxplot compares the computed enrichments to the ones obtained for randomly assigned reviewers to projects, showing that applicants are evenly distributed across projects. **d**. For each project, we compute the ratio between the proportion of applicant reviewers to the average proportion of applicant reviewers observed in the round. The boxplot compares the computed enrichments to the ones obtained for randomly assigned reviewers to projects, showing that applicants are evenly distributed across projects.

Such differences could be due to unfair grading, with reviewers from a certain category (applicants or non-applicants) grading more “randomly” than others. To analyse this effect, we need to look beyond average score into correlations. Indeed, two similar average scores could stem from highly different fine-grain score vectors. Imagine two reviewers grading 3 questions from 1 to 5. The first reviewer gives the grades 1, 2, and 5, while the second gives 5, 1, and 2. These reviews produce the same average score (2.67). However, their fine-grain structure is anti-correlated, with a Pearson correlation r = −0.69. In our context, we find that review scores are positively correlated, with a median Pearson correlation between their reviews of r = 0.28 across rounds (**Fig4d**), in line with previous observations in traditional funding schemes [35]. More importantly, we find no difference between applicants and non-applicants in their correlation with other project reviews (**Fig4c**). This indicates that the variability between grades within a review form are conserved across reviewer characteristics (see **Fig S7** and **Fig S9** for the other characteristics). As such, if applicants are uniformly distributed across projects, one will not expect a difference in the final rankings.

### A framework for iterative project implementations

In the JOGL implementation of the community review system, projects can apply to any number of rounds, irrespective of whether or not they have already successfully obtained funding in a previous round. We found 9 projects that applied to multiple rounds. On average, the relative performance of the projects in a grant round increases as a function of the number of participations (**Fig5a**). We find that this effect is explained by re-participation being associated with early success, with initially lower performing projects eventually dropping out (**Fig 5b-c**). As such, the multiple round scheme supports projects with a high initial potential in the long-term through repeated micro-funding allocations. We also note that in the case of 2 projects, re-participation after an initial failure allowed them to pass the acceptance threshold. This highlights how constructive feedback allows for a rapid improvement of a project and its successful re-application in the process.

**Figure 5.**
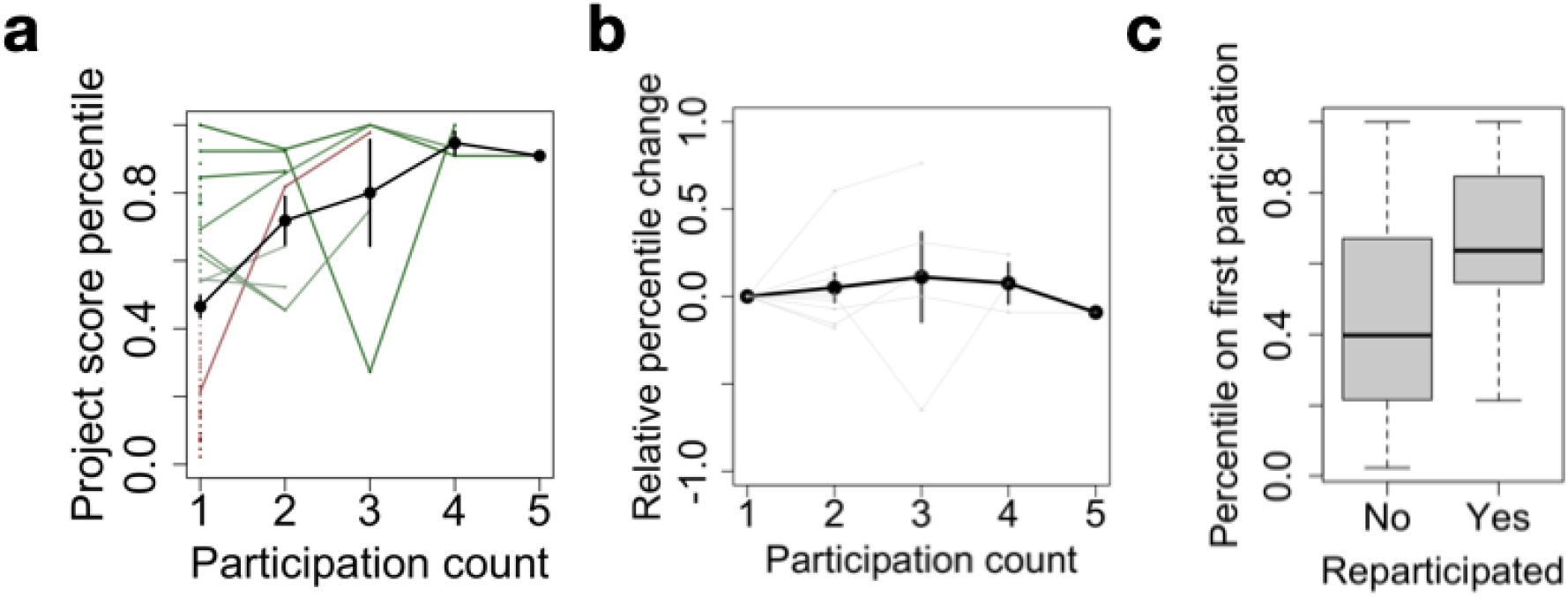
Multiple participations foster long-term project sustainability. **a** Project score percentile as a function of participation count. For each project, a score percentile is computed to quantify their relative rank within a specific application round, allowing to compare multiple projects across rounds. Participation count refers to the successive number of rounds a project has applied to. The black line denotes the average across projects, error bars represent standard error. Dots correspond to projects with only one participation, and lines to re-participating projects. Finally, the color gradient indicates relative score at first participation, from red (low) to green (high). **b** Same as a., after subtracting the percentile at first participation. **c** Score percentile at first participation as a function of whether or not a project has re-participated.

## Discussion

In this manuscript we describe the “community review” method for the identification of novel, feasible and potentially impactful projects within two communities of open innovation: Helpful Engineering and OpenCovid19. This process was leveraged for the attribution of volunteers as well as micro-grants to projects over a year, in an agile and iterative framework.

Key to the system is the requirement of applicants to take part in the reviewing process, ensuring its scalability. As such, the number of reviews is proportional to the number of projects applying (Fig 2), with a fast median process duration of 10 days. This requirement comes at a risk, since applicants might be negatively biased towards other projects they are competing against. Accordingly, we found that applicants consistently give a lower score to projects when compared to non-applicants (−0.52 points). This bias cannot be explained solely by the lower expertise of applicants towards the randomly assigned projects. Indeed, we found that self-reported expertise has only a limited impact on the final score (**Fig 4c**). The effect is most stringent for rare cases of self-reported expertise of 1 and 2 out of 5, suggesting that a threshold of 3 might be implemented to remove non-expert bias. It is on the other hand possible that non-applicants are positively biased towards projects from which they might have personally been invited to review. We however noted no such report in the conflict of interest question in the review form.

Despite these biases, we found that applicants and non-applicants have a similar behaviour when grading questions in the form, with a stable Pearson correlation between their reviews of r = 0.28 (**Fig4/Fig S8**). This is slightly higher than the correlation of 0.2 observed in an analysis of the ESRC’s existing peer review metrics (20), suggesting comparable outcomes when compared to existing institutional methods. The similarity of their correlation profiles means that such biases contribute a similar “noise” to the system: they might change the overall average scores, but not their ranking as long as applicants are well distributed across projects. Accordingly, we found that the community review system is robust to the removal of reviewers, with an average ranking Spearman correlation of 0.7 in the extreme case of one reviewer per project.

Finally, we showed that some projects apply multiple times to the application rounds. While the number of such projects of this type is small (9 projects), we find that it had two benefits. First, we found two projects that re-applied after an unsuccessful application, allowing them to pass the acceptance threshold on the second application. This showcases the ability of the feedback system to benefit projects in constructively improving their application. Furthermore, we found that the number of applications of a project is strongly dependent on its performance on the first application. This means that the iterative process allows to select highly promising projects and sustain their implementation in the mid- to long-term. This is of particular importance when considering traditional hackathon systems, where promising projects are usually not supported over longer periods of time.

The speed and cost-efficiency of the community review process has allowed for a reactive response to the high-pressure environment created by the pandemic. This agility has meant that within the short time frame given, projects have been able to produce literature, methods and hardware and put them to use (21–26). Overall, the community review system allows for a rapid, agile, iterative, distributed and scalable review process for volunteer action and micro-grant attribution. It is particularly suited for open research and innovation communities collaborating in a decentralized manner and looking for ways to distribute common resources fairly and swiftly. Finally, community review offers a robust alternative to institutional frameworks for building trust within a network and paves the way for the installation of community-driven decentralized laboratories.

## Supporting information

Sankey data input Figure 1c

Review form JOGL 1

Peer Review Form JOGL 2

Peer Review Form JOGL 3

Review form JOGL 4

Review form JOGL 5

Helpful Engineering Review Form

## Author Contributions

Conceptualisation of the micro grant allocation by TEL. Design of the grant review flows and community review by TEL and CLBG. Design of the peer review forms by TEL, CLBG, LB and MS. Iterative improvement of the method by TEL, CLBG and MS. Administration of community review by CLBG, TEL and EL. OpenCOVID19 initiative community coordination by TEL, CM and CLBG. Technical implementation within the JOGL platform by LB and LH. Initial grant dissemination by TEL, CLBG and EL with a special thanks to all reviewers. Data cleaning, processing and analysis of reviews by AV, CLBG and MS. Writing and figure creation by CLBG, BGT, TEL and MS.

## Supplementary Information

**Raw data of anonymised reviewers available on request**

**Raw reviewer forms in PDF attached on paper submission**

**Supplementary Project type and expertise Conversion tables 1 and 2 attached as a .csv**

## Supplementary Figures

**Figure S1a-c.**
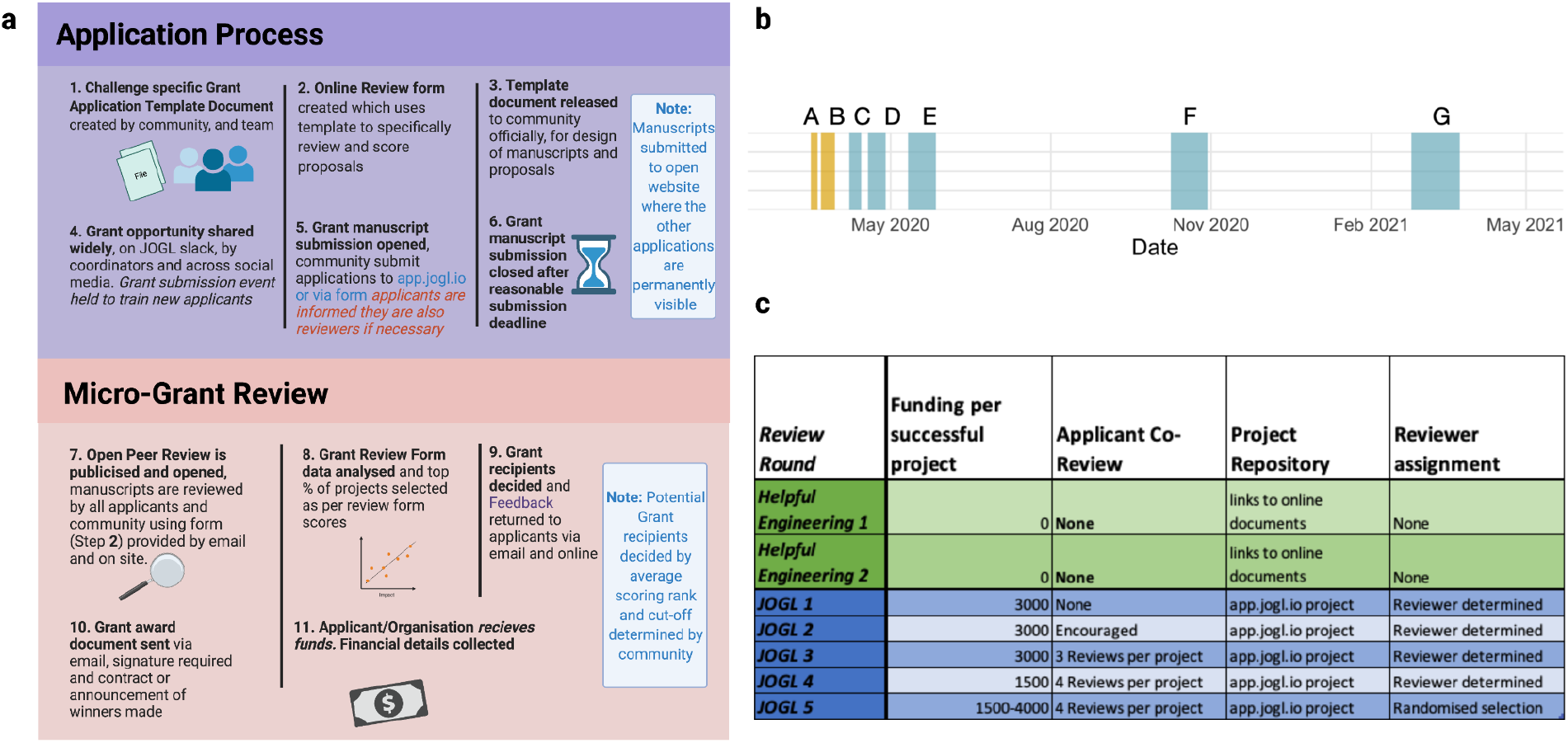
Methodology of review across programme project review rounds. **a**. Stages of the open peer review process and its improvement over time by applicant feedback. The online review forms and templates are found in supplementary data. Community refers to either the open voluntary community of Helpful Engineering, a COVID related charity, or JOGL, an online collaborative space for scientists and an already existing NGO, which based its peer review and grant allocation for COVID solutions on that of Helpful Engineering. **b** Time periods of each round **bA** 4 days -HE 1, **bB** 8 days HE 2, **bC** 7 Days - JOGL 1, **bD** 10 days - JOGL 2, **bE** 16 days - JOGL 3, **bF** 21 days - JOGL 4, **bG** 28 days JOGL 5, **c**. As the crisis developed, so did the purpose of peer review, from initially member allocation to funding allocation as projects matured. The administration and requirements for applicants changes over time as illustrated. HE= Helpful Engineering charity founded to produce open source designs for PPE and solutions to COVID, JOGL= Just One Giant Lab.

**Figure S2.**
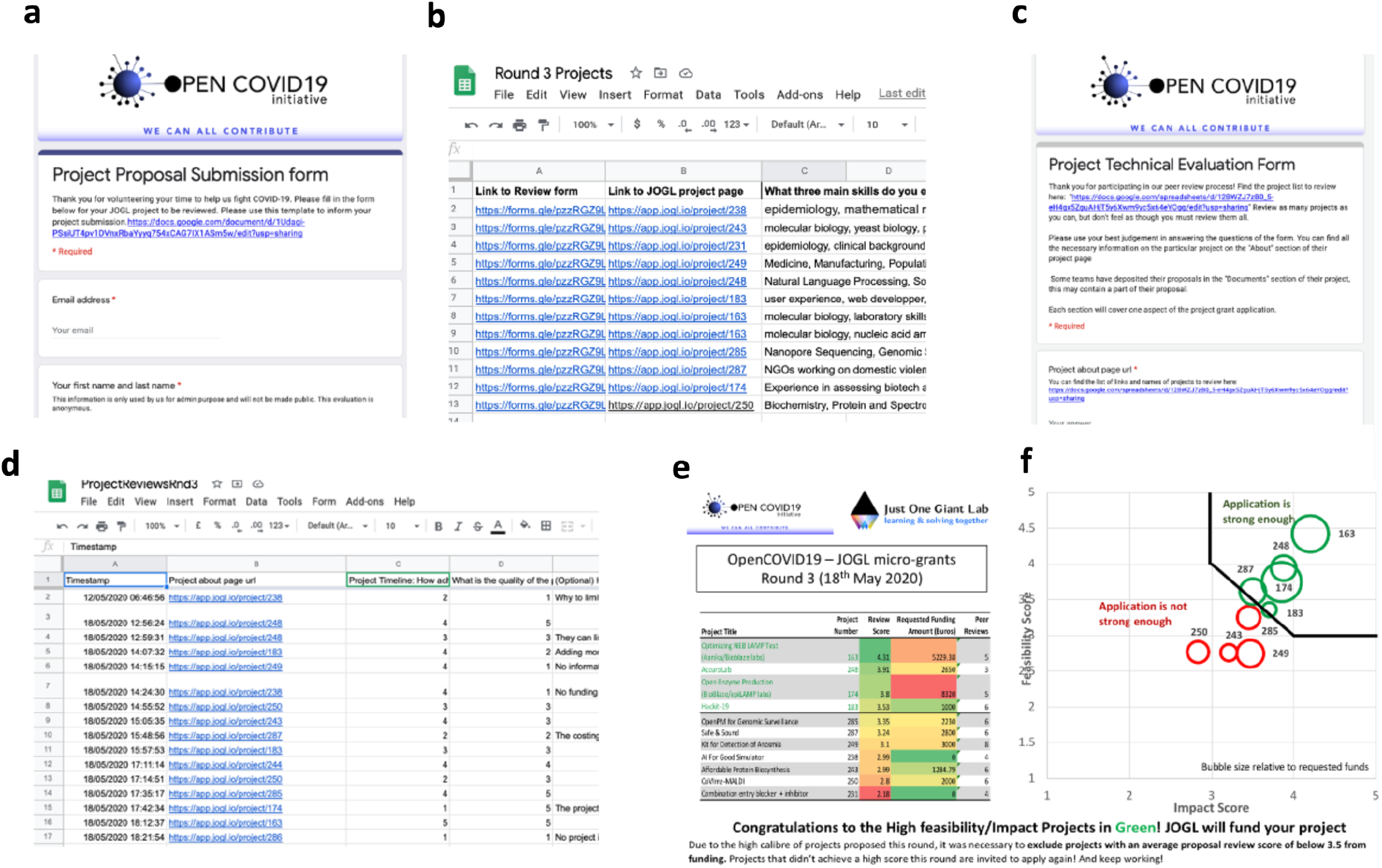
Process for online review using open tools. **a**. Submission of project proposals via a public google form. **b**. Open repository of proposals deposited to https://app.jogl.io/ link available in C. **c**. Evaluation via a Google form emailed to applicants and shared on available channels. **d**. Data collection of reviewer feedback. **e**. Public display of outcomes and scores, to assign grantees.

**Figure S3.**
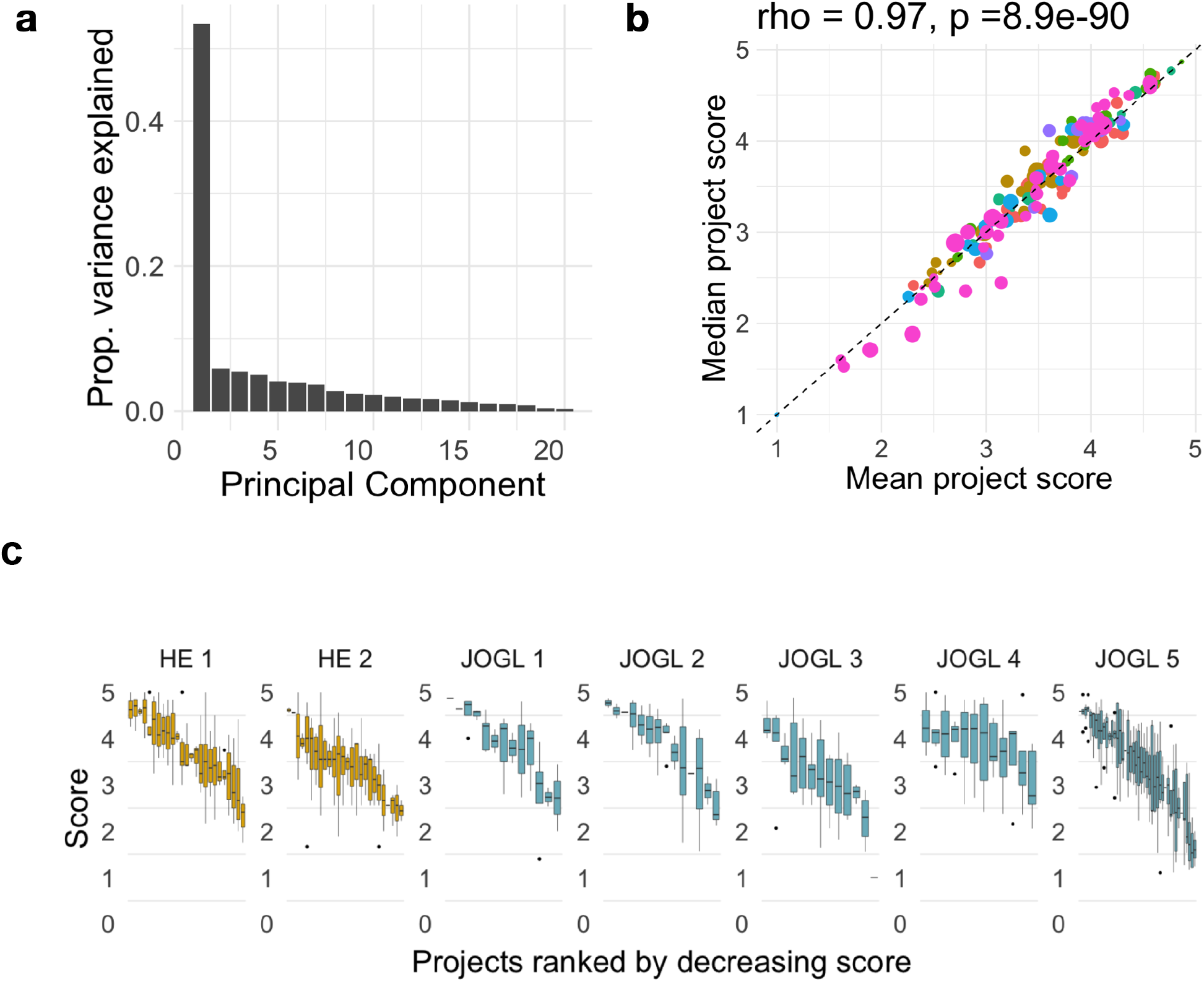
Project score variation. **a**. Principle component 1 dominance, indicating the importance of the explained variance in PC1 Figure 3b. **b**. Variance of projects, mean and median project scores, indicating mean or median as reliable scoring metrics. **c**. Scoring pattern of reviews to projects as boxplots across rounds.

**Figure S4.**
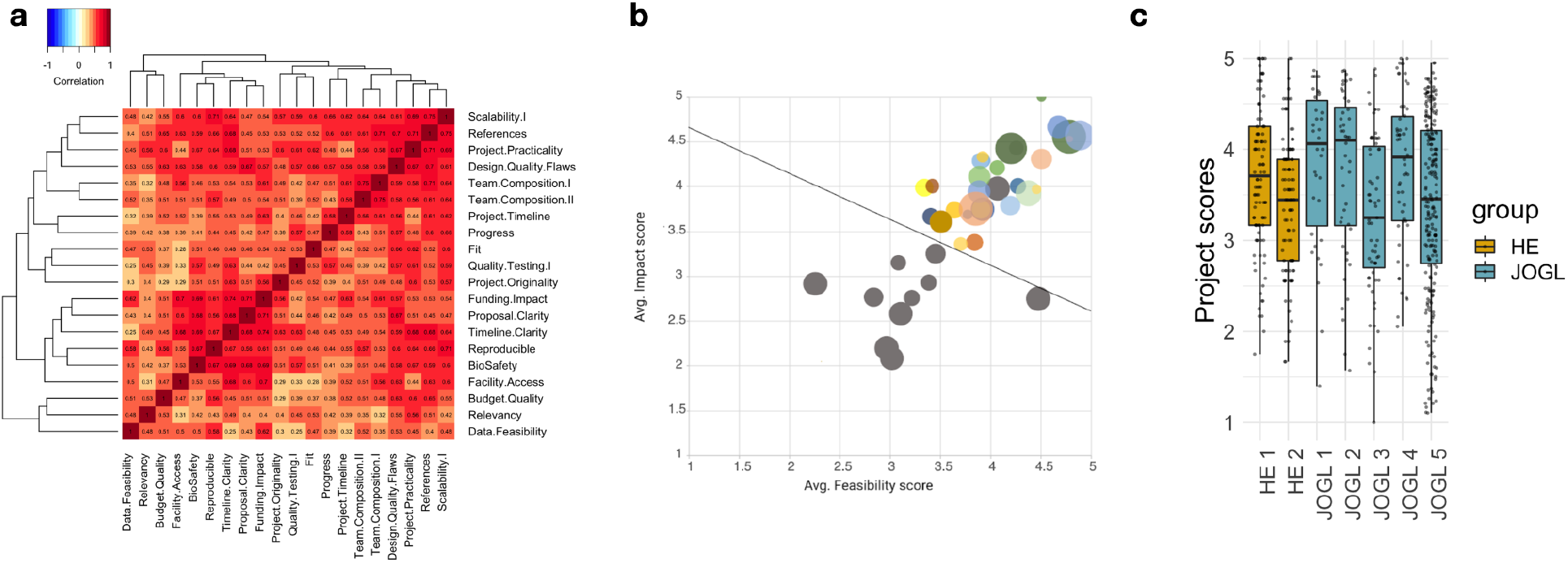
Correlation between question scores. **a** Heatmap showing the correlation matrix between question scores across reviews for JOGL round 4 (see original matrix in Figure 3a). Questions are overall strongly correlated with one another, as can be seen from the PCA in Fig3. **b** Projects from JOGL rounds 1-4 plotted as a function of their impact and feasibility scores. These correspond to two subset of questions from the form. As suggested by the correlation analysis, we find a strong correlation between the two quantities, indicating that the average score is a good proxy of project quality for most projects. We note that the projection into the feasibility/impact space allows to quickly visualise projects that are strong in one but not the other dimension, giving more granularity for the decision making process. **c** Average score distribution across review rounds.

**Figure S5.**
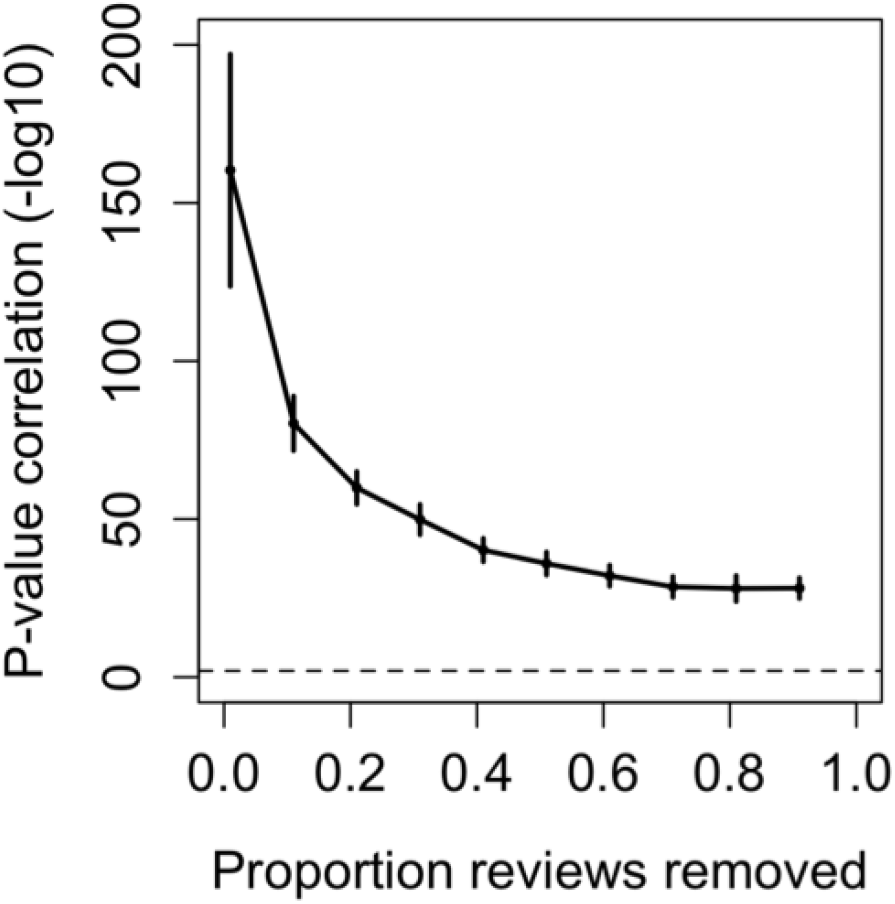
Significance of the correlations from Figure 3d (function cor.test in R)

**Figure S6.**
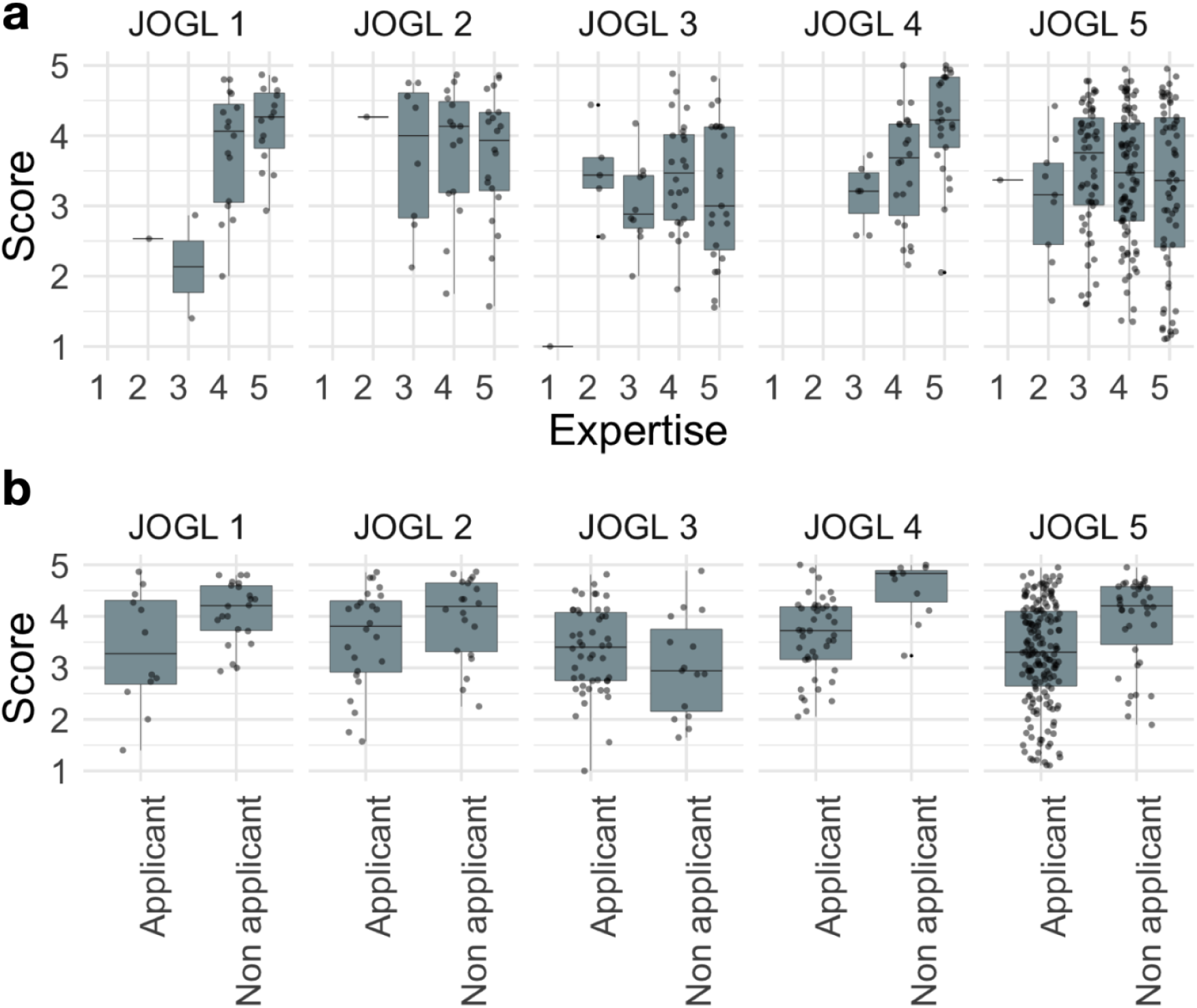
Same as Figure 5, per round.

**Figure S7.**
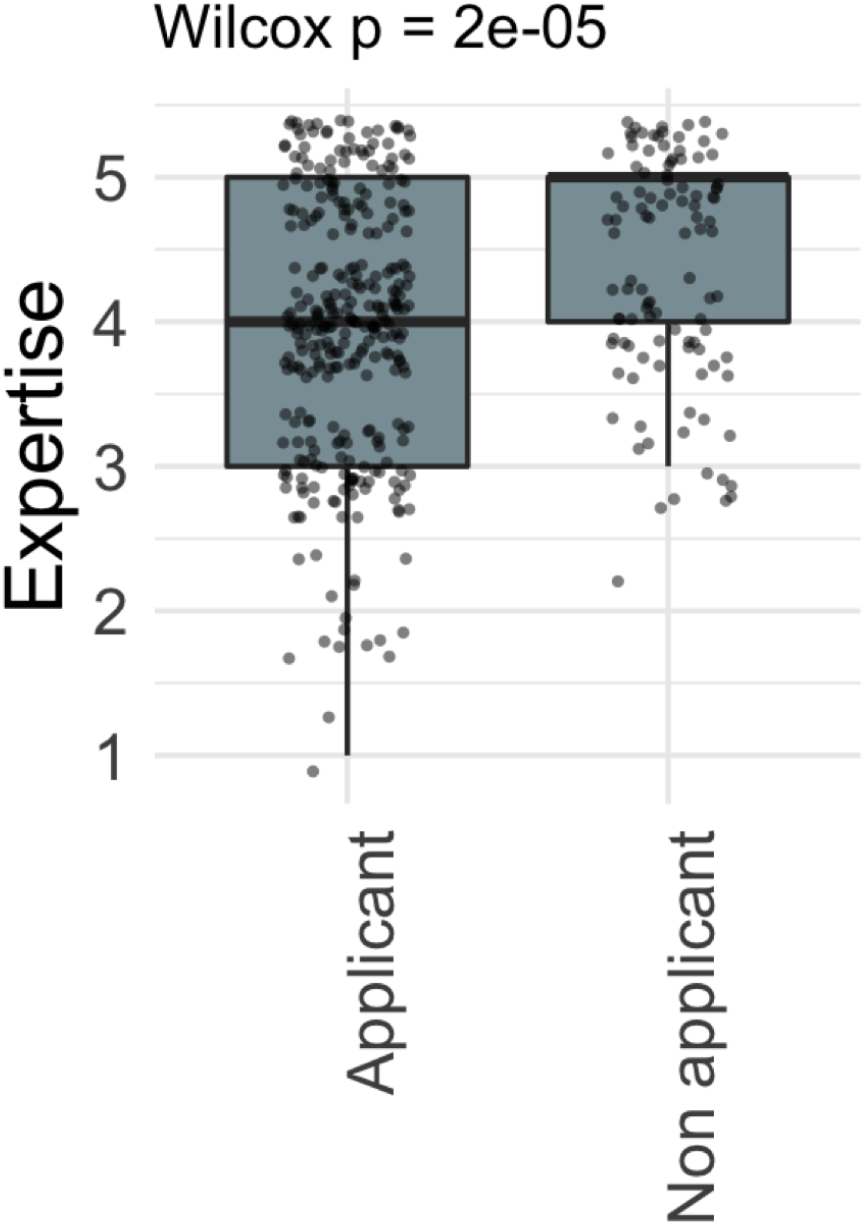
Expertise vs applicant status. We find that non-applicants are significantly more likely to be experts than applicants. This suggests that external reviewers who are volunteering to review do so because of their expertise towards the topic, while applicants are driven by the necessity to review other projects for their own projects to be reviewed. This is not found when JOGL staff members are removed **Fig S10**

**Figure S8.**
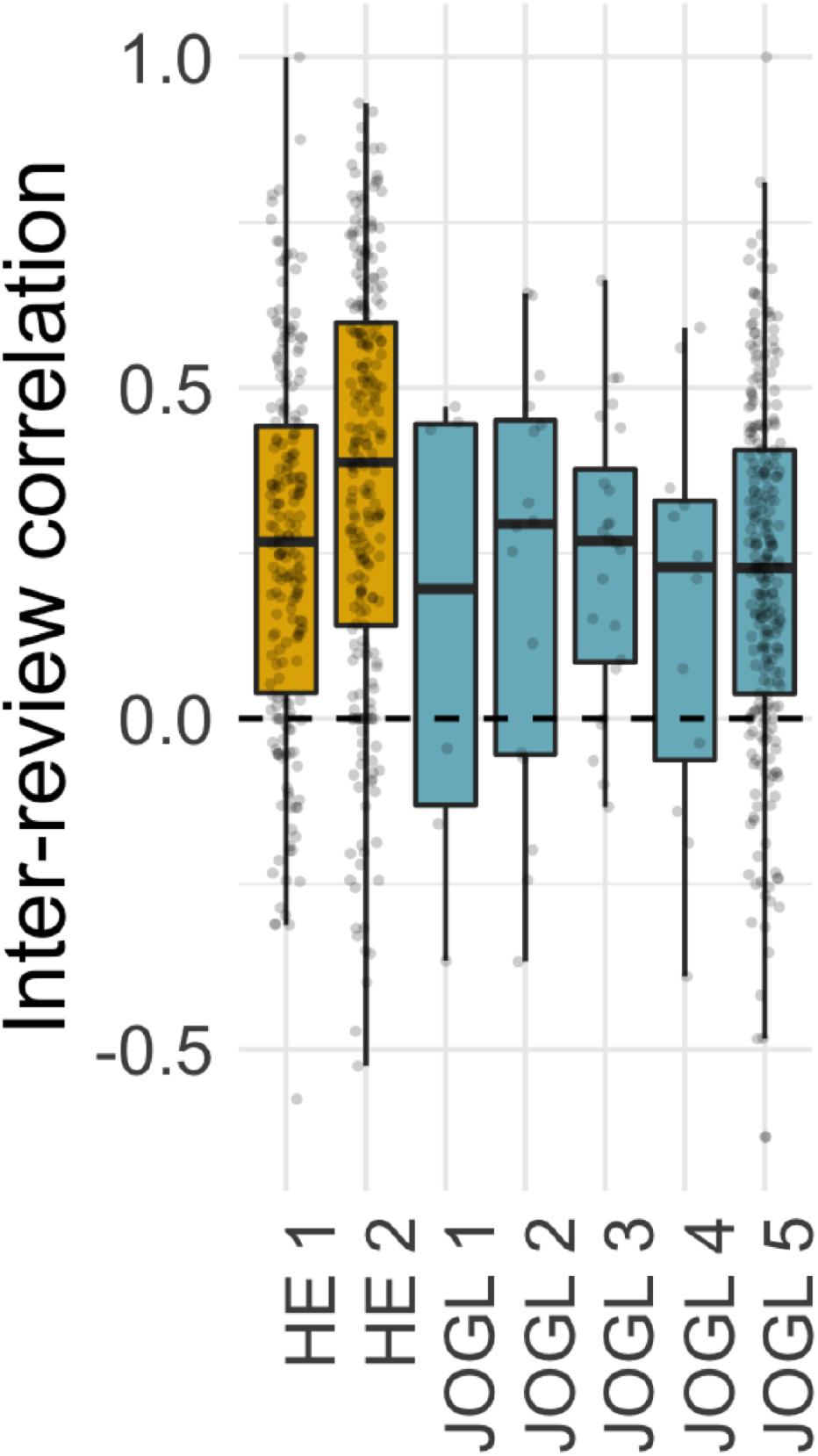
Inter-review correlation per round. Round correlations indicate a reliable correlation across rounds independent of scale. Correlation refers to agreement on scoring patterns between reviewers.

**Figure S9.**
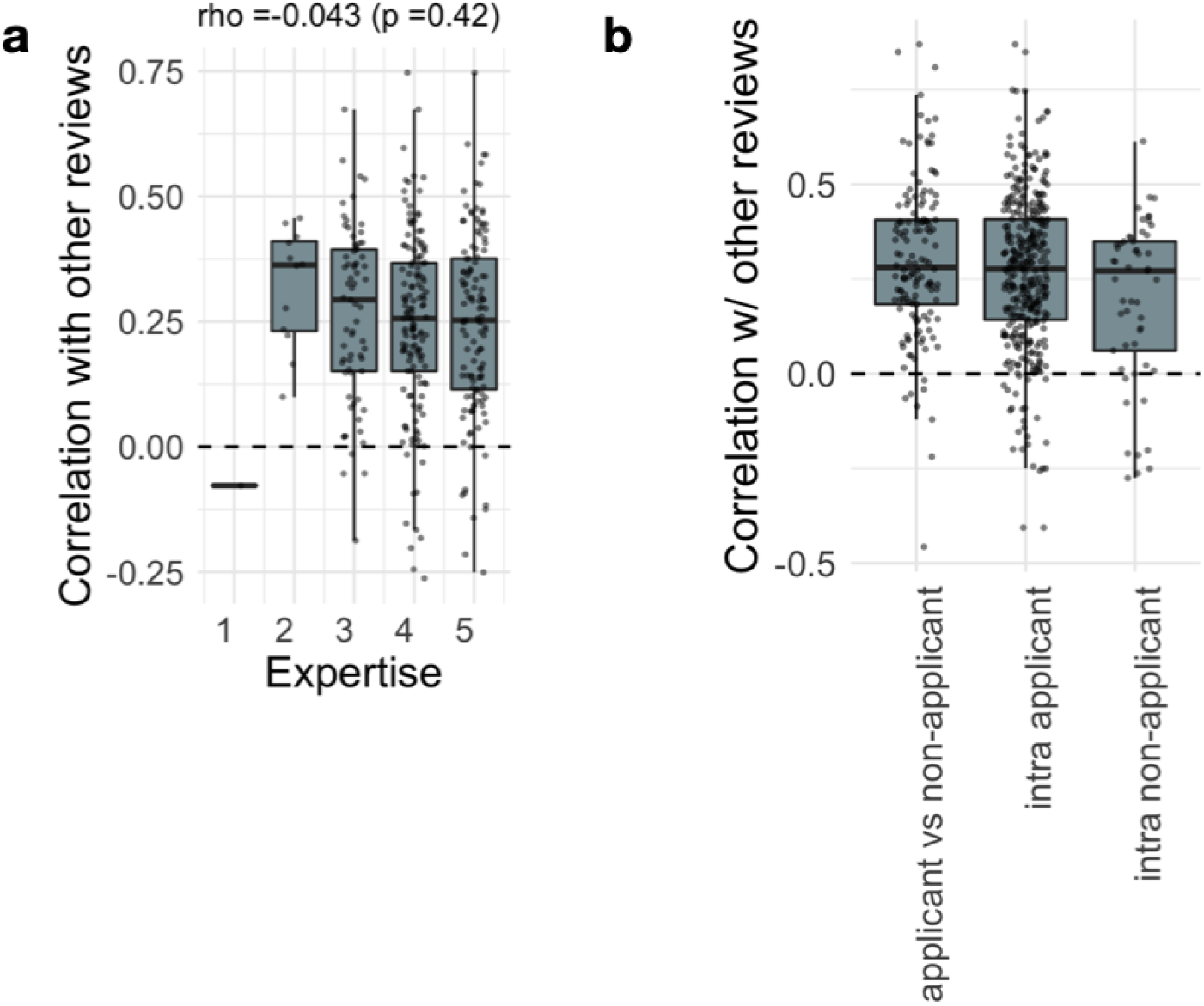
**a-b** Same as Figure 6b, for expertise level and self-reviewing status. We find a small effect where self-reviewers are less correlated with other reviewers compared to independent reviewers, but the numbers are too small to conclude a significant result. **c** We compute the correlation between project reviews for specific pairs of reviewers based on their application status. We find that non-applicants have a smaller correlation with other non-applicants compared to other pairs (Mann-Whitney U test, p = 8.6e-3 compared to “applicant vs non-applicant” and p = 7.3e-2 when compared with intra-applicant).

**Figure S10.**
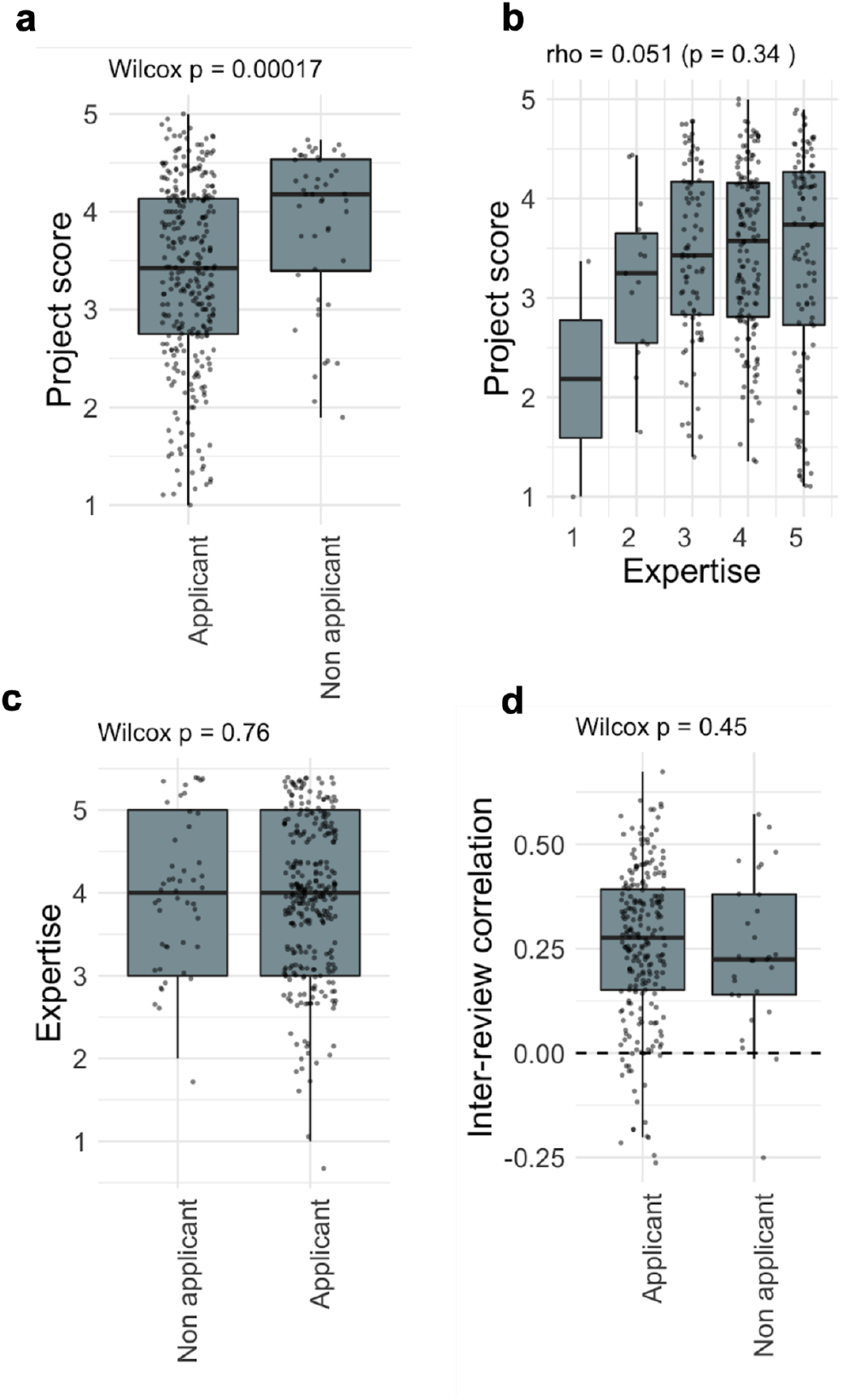
Reviewer correlations after removal of reviewers who are JOGL staff members from the analysis. Project scores, expertise distributions, and interview correlations remained the same after removal of JOGL staff members who partook in the review process as part of the community. **a**. Project score vs application status t **b**. Project score vs expertise of reviewer **c**. Expertise of applicants vs non applicants **d**. Inter-review correlations applicant vs non applicant

**Supplementary Table 1.** OLS model of review score as a function of expertise (1-5) and applicant (0: non-applicant, 1: applicant).

| Residuals |  |  |  |  |
| --- | --- | --- | --- | --- |
| Min | 1Q | Median | 3Q | Max |
| -2.2926 | -0.6239 | 0.1332 | 0.7539 | 1.6428 |
| Coefficients |  |  |  |  |

|  | **Estimate** | **Std. Error** | **t value** | **Pr(>t)** |
| --- | --- | --- | --- | --- |
| (Intercept) | 3.7155 | 0.25158 | 14.769 | <2e-16 |
| expertise | 0.04065 | 0.05388 | 0.754 | 0.451 |
| applicant | -0.52094 | 0.10701 | -4.868 | 1.61e-06 |

## Notes

### Competing Interest Statement

The authors have declared no competing interest.

### Summary of Updates

Author contributions updated, and minor grammatical additions.

