## Supplementary material for "Community review: a robust and scalable selection system for resource allocation within open science and innovation communities": Review form JOGL 1

### Project Technical Evaluation Form

Thank you for participating in our peer review process! Find the project list to review here: "https://docs.google.com/spreadsheets/d/1CQATJl-tyllZyqfHpJqHHe03wxIOEkKPKEdWoCGu-OA/edit?usp=sharing" Review as many projects as you can, but don't feel as though you must review them all.

Please use your best judgement in answering the questions of the form. You can find all the necessary information on the particular project on the "About" section of their project page, the project must have followed the template guidelines written here: https://docs.google.com/document/d/1Udagi-PSsiUT4pv1DVnxRbaYyyq754xCAG7IX1ASm5w/edit. Some teams have deposited their proposals in the "Documents" section of their project, this may contain a part of their proposal.

Each section will cover one aspect of the project grant application.

* Required

1. Project about page url *

You can find the list of links and names of projects to review here: "https://docs.google.com/spreadsheets/d/1CQATJl-tyllZyqfHpJqHHe03wxIOEkKPKEdWoCGu-OA/edit?usp=sharing"

#### Feasibility

2. Project Timeline: How achievable is this project within 3 months? *

1. The project is not achievable | 3. The project is achievable, but will take a great effort by all involved and more participants | 5. The project is very achievable with little extra input required.

*Mark only one oval.*

| 1 | 2 | 3 | 4 | 5 |
|---|---|---|---|---|
| ⬭ | ⬭ | ⬭ | ⬭ | ⬭ |

3. Rough Final cost of project in euros *

*Mark only one oval.*

- ⬭ 0
- ⬭ 50
- ⬭ 100
- ⬭ 500
- ⬭ 1000
- ⬭ 2500-5000
- ⬭ 5000+

4. Do you agree with the estimated cost of the project? *

1. Project underestimates cost | 3. Project has accurate costing | 5. Project overestimates cost

*Mark only one oval.*

| 1 | 2 | 3 | 4 | 5 |
|---|---|---|---|---|
| ⬭ | ⬭ | ⬭ | ⬭ | ⬭ |

5. (Optional) How could the project cost be improved/reduced? Please provide any details of contacts if applicable.

6. Is the project's current team composition sufficient to achieve the projects goal? *

1. Project underestimates the team needed, and doesn't have enough members| 3. The team composition seems "just" sufficient to be achievable | 5. The project has many active team members, and enough to achieve its goal well.

*Mark only one oval.*

| 1 | 2 | 3 | 4 | 5 |
|---|---|---|---|---|
| ⬭ | ⬭ | ⬭ | ⬭ | ⬭ |

7. Material Sources *

*Mark only one oval.*

- Retail stores
- Online Retail
- Industrial
- Medical grade
- Laboratory only
- Computational

8. Does the project have access to the materials/ facilities required to complete the project?

1. The project does not have access to the materials or facilities required| 3. The project has access to materials/facilities, however not enough to fully scale the project | 5. The project has enough facility access to be completed to fruition.

*Mark only one oval.*

| 1 | 2 | 3 | 4 | 5 |
|---|---|---|---|---|
| ⬭ | ⬭ | ⬭ | ⬭ | ⬭ |

9. How clear is the project proposal? *

1. Unclear and/or irrelevant | 3. Clear but relevant points are missing | 5. Very clear and relevant

*Mark only one oval.*

| 1 | 2 | 3 | 4 | 5 |
|---|---|---|---|---|
| ⬭ | ⬭ | ⬭ | ⬭ | ⬭ |

10. Project design: What is the project design quality? Are there flaws? *

1. Many problems with project design | 3. Design seems feasible but it's inefficient | 5. Brilliant Design, well documented.

*Mark only one oval.*

| 1 | 2 | 3 | 4 | 5 |
|---|---|---|---|---|
| ⬭ | ⬭ | ⬭ | ⬭ | ⬭ |

11. Which category would you say the project falls under? *

*Mark only one oval.*

- ⬭ Medical Hardware Design/Adaptation
- ⬭ Self Screening (Software)
- ⬭ Other Software
- ⬭ Diagnostic technology
- ⬭ Data Analysis/Modelling project
- ⬭ PPE
- ⬭ Other: ____________________

12. Clarity and relevance of the project timeline and it's needs for future (major tasks, milestones) *

1. Unclear and irrelevant | 3. Clear but relevant points are missing | 5. Very clear and relevant

*Mark only one oval.*

| 1 | 2 | 3 | 4 | 5 |
|---|---|---|---|---|
| ⬭ | ⬭ | ⬭ | ⬭ | ⬭ |

13. If this project is biological in nature, do they address the biosafety and regulations requirements to address? (if applicable)

1. Not at all | 3. Aware but unclear integration | 5. Actively engaged and addressed these regulations

*Mark only one oval.*

| 1 | 2 | 3 | 4 | 5 |
|---|---|---|---|---|
| ⬭ | ⬭ | ⬭ | ⬭ | ⬭ |

14. If it is a data driven project how feasible is the data collection for this project ? (Really Hard / The data exists already) (Data analytics projects only)

1. Not at all | 3. Aware but unclear integration | 5. Actively engaged and addressed these regulations

*Mark only one oval.*

| 1 | 2 | 3 | 4 | 5 |
|---|---|---|---|---|
| ⬭ | ⬭ | ⬭ | ⬭ | ⬭ |

15. What is the project's state of progress? *

1. No results yet | 3. Some results but with a promising development plan | 5. Working proof of concept

*Mark only one oval.*

| 1 | 2 | 3 | 4 | 5 |
|---|---|---|---|---|
| ⬭ | ⬭ | ⬭ | ⬭ | ⬭ |

#### Project Type

16. Is this project a product or data analysis/model *

*Mark only one oval.*

- ⬭ A data analysis project/model *Skip to question 23*
- ⬭ A Software/Hardware product/New Method Eg: Diagnostic test

#### Product Impact

17. Fit between the project's approach/methodology and the problem they have stated to resolve *

1. Inappropriate | 3. Appropriate | 5.Very appropriate

*Mark only one oval.*

| 1 | 2 | 3 | 4 | 5 |
|---|---|---|---|---|
| ⬭ | ⬭ | ⬭ | ⬭ | ⬭ |

18. How relevant is the project to the COVID-19 crisis and alleviation of suffering? *

1. Not at all | 3. Moderately | 5. Essential

*Mark only one oval.*

| 1 | 2 | 3 | 4 | 5 |
|---|---|---|---|---|
| ⬭ | ⬭ | ⬭ | ⬭ | ⬭ |

19. The scalability of the project in the long term, taking manufacturing into consideration *

1. No sustainability model and no scaling potential | 3. Sustainable and scalable but still unstructured plan | 5. Already applying a sustainable plan & good scalability

*Mark only one oval.*

| 1 | 2 | 3 | 4 | 5 |
|---|---|---|---|---|
| ⬭ | ⬭ | ⬭ | ⬭ | ⬭ |

20. How well has the project addressed concerns such as quality testing? *

1. Not well | 3. Clear but not entirely appropriate | 5. Very clear and relevant

*Mark only one oval.*

| 1 | 2 | 3 | 4 | 5 |
|---|---|---|---|---|
| ⬭ | ⬭ | ⬭ | ⬭ | ⬭ |

21. The project's dissemination strategy (quality of documentation for goals, results, methods and needs; open access model, outreach) is this/will this project be easily reproducible? *

1. Irreproducible and poorly documented 3. Good documentation but with few weaknesses 5. Easily reproducible & very well communicated

*Mark only one oval.*

| 1 | 2 | 3 | 4 | 5 |
|---|---|---|---|---|
| ⬭ | ⬭ | ⬭ | ⬭ | ⬭ |

22. Project Originality *

How much is this app/device needed? Are there already apps or devices that can do this job in an open source way? 1. The project is unoriginal and done before | 3. It has been done before, but not in an open source way | 5. The project is completely original as far as the reviewer is aware.

*Mark only one oval.*

| 1 | 2 | 3 | 4 | 5 |
|---|---|---|---|---|
| ⬭ | ⬭ | ⬭ | ⬭ | ⬭ |

*Skip to question 30*

#### Data Analysis project Impact

23. Fit between the project's approach/methodology and the problem they have stated to resolve *

1. Inappropriate | 3. Appropriate | 5.Very appropriate

*Mark only one oval.*

| 1 | 2 | 3 | 4 | 5 |
|---|---|---|---|---|
| ⬭ | ⬭ | ⬭ | ⬭ | ⬭ |

24. How relevant will the result of this project be for policy makers? *

1. Not at all | 3. Moderately | 5. Essential

*Mark only one oval.*

| 1 | 2 | 3 | 4 | 5 |
|---|---|---|---|---|
| ⬭ | ⬭ | ⬭ | ⬭ | ⬭ |

25. How relevant will the result of this project be for the public? *

1. Not at all | 3. Moderately | 5. Essential

*Mark only one oval.*

| 1 | 2 | 3 | 4 | 5 |
|---|---|---|---|---|
| ◯ | ◯ | ◯ | ◯ | ◯ |

26. The project's dissemination strategy (quality of documentation for goals, results, methods and needs; open access model, outreach) how reproducible does this project seem? *

1. Irreproducible and poorly documented 3. Good documentation but with few weaknesses 5. Easily reproducible & very well communicated

*Mark only one oval.*

| 1 | 2 | 3 | 4 | 5 |
|---|---|---|---|---|
| ◯ | ◯ | ◯ | ◯ | ◯ |

27. How accessible will the result of the project be to the general public? *

*Mark only one oval.*

| | 1 | 2 | 3 | 4 | 5 | |
|---|---|---|---|---|---|---|
| Unaccessible | ◯ | ◯ | ◯ | ◯ | ◯ | Easily accessible and distributed on media |

28. How accessible will the result of the project be to policy makers? *

*Mark only one oval.*

| | 1 | 2 | 3 | 4 | 5 | |
|---|---|---|---|---|---|---|
| Unaccessible | ◯ | ◯ | ◯ | ◯ | ◯ | Easily accessible and distributed to policy makers specifically |

29. Project Originality *

How much is this analysis needed/unique? Are there already open source analysis's pipelines out there that can do this? 1. Unoriginal and done before/not needed | 3.Done before but not relevant to this crisis | 5. The project is original as far as the reviewer is aware.

*Mark only one oval.*

| 1 | 2 | 3 | 4 | 5 |
|---|---|---|---|---|
| ◯ | ◯ | ◯ | ◯ | ◯ |

#### Additional comments and reviewer self-evaluation

You may provide below additional comments addressed to the team leaders of the project, and private comments to the admin team.

Please, also assess your expertise in the fields relevant to evaluate the project.

30. [To the team leaders] Other general constructive comments about the project *

31. [To admins] Private comment - anything you would like to inform the admin team of, privately.

32. What is your expertise that is relevant to evaluate this project? *

Please provide justification of your expertise if possible.

33. How would you assess your own abilities to review this project? *

*Mark only one oval.*

| | 1 | 2 | 3 | 4 | 5 | |
|---|---|---|---|---|---|---|
| I cannot be trusted to assess this project, and did not fully grasp the project. | ◯ | ◯ | ◯ | ◯ | ◯ | I can fully assess this project, based on my pr |

34. Are you part of the project team that you are currently assessing? *

*Mark only one oval.*

◯ Yes
◯ No

35. Are you part of a project team within the OpenCOVID19 initiative? *

*Mark only one oval.*

◯ Yes
◯ No

36. Would you like us to include you in our "Experts community" and provide similar reviews to projects when you have time in the future? *

*Mark only one oval.*

- Yes
- No

37. What is your JOGL username ?(if applicable)

_______________________________

38. What is your Slack Handle on the OpenCovid19 slack?(if applicable)

_______________________________

This content is neither created nor endorsed by Google.

Google Forms
