## Supplementary material for "Community review: a robust and scalable selection system for resource allocation within open science and innovation communities": Peer Review Form JOGL 3

### Project Technical Evaluation Form

Thank you for participating in our peer review process! Find the project list to review here: "https://docs.google.com/spreadsheets/d/12BWZJ7zB0_5-eH4gx5ZguAHjT5y6Xwm9yc5x64eYOgg/edit?usp=sharing" Review as many projects as you can, but don't feel as though you must review them all.

Please use your best judgement in answering the questions of the form. You can find all the necessary information on the particular project on the "About" section of their project page

Some teams have deposited their proposals in the "Documents" section of their project, this may contain a part of their proposal.

Each section will cover one aspect of the project grant application.

* Required

1. Project url *

You can find the list of links and names of projects to review here: https://docs.google.com/spreadsheets/d/12BWZJ7zB0_5-eH4gx5ZguAHjT5y6Xwm9yc5x64eYOgg/edit?usp=sharing

#### Feasibility

2. Project Timeline: How achievable is this project within 3 months? *

1. The project is not achievable | 3. The project is achievable, but will take a great effort by all involved and more participants | 5. The project is very achievable with little extra input required.

*Mark only one oval.*

| 1 | 2 | 3 | 4 | 5 |
|---|---|---|---|---|
| ◯ | ◯ | ◯ | ◯ | ◯ |

3. What is the quality of the project budgeting? *

1. Low Quality/Non existent. 3. Costing exists and has specific costs allocated. 5. Costing is fully transparent and documented.

*Mark only one oval.*

| 1 | 2 | 3 | 4 | 5 |
|---|---|---|---|---|
| ◯ | ◯ | ◯ | ◯ | ◯ |

4. (Optional) How could the project cost be improved/reduced? Please provide any details of contacts if applicable.

5. Is the project's current team composition sufficient to achieve the projects goal? *

1. Project underestimates the team needed, and doesn't have enough members| 3. The team composition seems "just" sufficient to be achievable | 5. The project has many active team members, and enough to achieve its goal well.

*Mark only one oval.*

| 1 | 2 | 3 | 4 | 5 |
|---|---|---|---|---|
| ⬭ | ⬭ | ⬭ | ⬭ | ⬭ |

6. Does the project have access to the materials/ facilities required to complete the project?

1. The project does not have access to the materials or facilities required| 3. The project has access to materials/facilities, however not enough to fully scale the project | 5. The project has enough facility access to be completed to fruition.

*Mark only one oval.*

| 1 | 2 | 3 | 4 | 5 |
|---|---|---|---|---|
| ⬭ | ⬭ | ⬭ | ⬭ | ⬭ |

7. How clear is the project proposal? *

1. Unclear and/or irrelevant | 3. Clear but relevant points are missing | 5. Very clear and relevant

*Mark only one oval.*

| 1 | 2 | 3 | 4 | 5 |
|---|---|---|---|---|
| ⬭ | ⬭ | ⬭ | ⬭ | ⬭ |

8. Project design: What is the project design quality? Are there flaws? *

1. Many problems with project design | 3. Design seems feasible but it's inefficient | 5. Brilliant Design, well documented.

*Mark only one oval.*

| 1 | 2 | 3 | 4 | 5 |
|---|---|---|---|---|
| ⬭ | ⬭ | ⬭ | ⬭ | ⬭ |

9. Which category would you say the project falls under? *

*Mark only one oval.*

- ⬭ Medical Hardware Design/Adaptation
- ⬭ Self Screening (Software)
- ⬭ Other Software
- ⬭ Diagnostic technology
- ⬭ Data Analysis/Modelling project
- ⬭ PPE
- ⬭ Other: ______________________

10. Clarity and relevance of the project timeline and it's needs for future (major tasks, milestones) *

1. Unclear and irrelevant | 3. Clear but relevant points are missing | 5. Very clear and relevant

*Mark only one oval.*

| 1 | 2 | 3 | 4 | 5 |
|---|---|---|---|---|
| ⬭ | ⬭ | ⬭ | ⬭ | ⬭ |

11. If this project is biological in nature, do they address the biosafety and regulations requirements to address? (if applicable)

1. Not at all | 3. Aware but unclear integration | 5. Actively engaged and addressed these regulations

*Mark only one oval.*

| 1 | 2 | 3 | 4 | 5 |
|---|---|---|---|---|
| ⬭ | ⬭ | ⬭ | ⬭ | ⬭ |

12. If it is a data driven project how feasible is the data collection for this project ? (Really Hard / The data exists already) (Data analytics projects only)

1. Not at all | 3. Aware but unclear integration | 5. Actively engaged and addressed these regulations

*Mark only one oval.*

| 1 | 2 | 3 | 4 | 5 |
|---|---|---|---|---|
| ⬭ | ⬭ | ⬭ | ⬭ | ⬭ |

13. What is the project's state of progress? *

1. No results yet | 3. Some results but with a promising development plan | 5. Working proof of concept

*Mark only one oval.*

| 1 | 2 | 3 | 4 | 5 |
|---|---|---|---|---|
| ⬭ | ⬭ | ⬭ | ⬭ | ⬭ |

#### Project Type

14. Is this project a product or data analysis/model *

*Mark only one oval.*

⬭ A data analysis project/model *Skip to question 22*

⬭ A Software/Hardware product/New Method Eg: Diagnostic test

*Mark only one oval.*

- Yes
- No

38. What is your JOGL username ?(if applicable)

_______________________________________________

39. What is your Slack Handle on the OpenCovid19 slack?(if applicable)

_______________________________________________

This content is neither created nor endorsed by Google.

Google Forms
