## Supplementary material for "Community review: a robust and scalable selection system for resource allocation within open science and innovation communities": Review form JOGL 4

### Project Technical Evaluation Form

Thank you for participating in our peer review process! Find the project list to review here: "https://docs.google.com/spreadsheets/d/1jodjvIpLY5rQ03y00sBBdP5yrKmZNJ_Fo86h5jNpUNg/edit?usp=sharing" Review as many projects as you can, but don't feel as though you must review them all.

Please use your best judgement in answering the questions of the form. You can find all the necessary information on the particular project on the "About" section of their project page

Some teams have deposited their proposals in the "Documents" section of their project, this may contain a part of their proposal.

Each section will cover one aspect of the project grant application.

* Required

1. Email *

2. Project URL Link *

You can find the list of links and names of projects to review here: https://docs.google.com/spreadsheets/d/1jodjvIpLY5rQ03y00sBBdP5yrKmZNJ_Fo86h5jNpUNg/edit?usp=sharing

#### Feasibility

3. Project Timeline: How achievable is this project within 3 months? *

1. The project is not achievable | 3. The project is achievable, but will take a great effort by all involved and more participants | 5. The project is very achievable with little extra input required.

*Mark only one oval.*

| 1 | 2 | 3 | 4 | 5 |
|---|---|---|---|---|
| ⬭ | ⬭ | ⬭ | ⬭ | ⬭ |

4. What is the quality of the project budgeting? *

1. Low Quality/Non existent. 3. Costing exists and has specific costs allocated. 5. Costing is fully transparent and documented.

*Mark only one oval.*

| 1 | 2 | 3 | 4 | 5 |
|---|---|---|---|---|
| ⬭ | ⬭ | ⬭ | ⬭ | ⬭ |

5. (Optional) How could the project cost be improved/reduced? Please provide any details of contacts if applicable.

6. Is the project's current team composition sufficient to achieve the projects goal? Part I *

1. Project underestimates the team needed, and doesn't have enough members| 3. The team composition seems "just" sufficient to be achievable | 5. The project has many active team members, and enough to achieve its goal well.

*Mark only one oval.*

| 1 | 2 | 3 | 4 | 5 |
|---|---|---|---|---|
| ◯ | ◯ | ◯ | ◯ | ◯ |

7. Is the project's current team composition sufficient to achieve the projects goal? Part II *

1. Project team members inexperienced, or no details of team given s| 3. The team expertise seem sufficient to achieve the goals set out by the project | 5. The project has multiple highly distinguished and or/experienced members who have a history of completing similar projects.

*Mark only one oval.*

| 1 | 2 | 3 | 4 | 5 |
|---|---|---|---|---|
| ◯ | ◯ | ◯ | ◯ | ◯ |

8. Does the project have access to the materials/ facilities required to complete the project?

1. The project does not have access to the materials or facilities required| 3. The project has access to materials/facilities, however not enough to fully scale the project | 5. The project has enough facility access to be completed to fruition.

*Mark only one oval.*

| 1 | 2 | 3 | 4 | 5 |
|---|---|---|---|---|
| ◯ | ◯ | ◯ | ◯ | ◯ |

9. How clear is the project proposal? *

1. Unclear and/or irrelevant | 3. Clear but relevant points are missing | 5. Very clear and relevant

*Mark only one oval.*

| 1 | 2 | 3 | 4 | 5 |
|---|---|---|---|---|
| ◯ | ◯ | ◯ | ◯ | ◯ |

10. Project design I: What is the project design quality? Are there flaws? *

1. Many problems with project design | 3. Design seems feasible but it's inefficient | 5. Brilliant Design, well documented.

*Mark only one oval.*

| 1 | 2 | 3 | 4 | 5 |
|---|---|---|---|---|
| ◯ | ◯ | ◯ | ◯ | ◯ |

11. Project design II: References and Prior Research, is there good foundation for this project? (Not applicable to some hardware of humanitarian projects)

1. No references or reference to published research or projects (if applicable) | 3. References poor in relevance, or their own preliminary data not shown (If applicable) | 5. Multiple relevant references, and/or preliminary data shown -indicating a good foundation.(If applicable)

*Mark only one oval.*

| 1 | 2 | 3 | 4 | 5 |
|---|---|---|---|---|
| ⬭ | ⬭ | ⬭ | ⬭ | ⬭ |

12. Which category would you say the project falls under? *

*Mark only one oval.*

- ⬭ Medical Hardware Design/Adaptation
- ⬭ Self Screening (Software)
- ⬭ Other Software
- ⬭ Diagnostic technology
- ⬭ Data Analysis/Modelling project
- ⬭ PPE
- ⬭ Other: ____________________

13. Clarity and relevance of the project timeline and it's needs for future (major tasks, milestones) *

1. Unclear and irrelevant | 3. Clear but relevant points are missing | 5. Very clear and relevant

*Mark only one oval.*

| 1 | 2 | 3 | 4 | 5 |
|---|---|---|---|---|
| ⬭ | ⬭ | ⬭ | ⬭ | ⬭ |

14. If this project is biological in nature, do they address the biosafety and regulations requirements to address? (if applicable)

1. Not at all | 3. Aware but unclear integration | 5. Actively engaged and addressed these regulations

*Mark only one oval.*

| 1 | 2 | 3 | 4 | 5 |
|---|---|---|---|---|
| ⬭ | ⬭ | ⬭ | ⬭ | ⬭ |

15. If it is a data driven project how feasible is the data collection for this project ? (Really Hard / The data exists already) (Data analytics projects only)

1. Not at all | 3. Aware but unclear integration | 5. Actively engaged and addressed these regulations

*Mark only one oval.*

| 1 | 2 | 3 | 4 | 5 |
|---|---|---|---|---|
| ⬭ | ⬭ | ⬭ | ⬭ | ⬭ |

16. What is the project's state of progress? *

1. No results yet | 3. Some results but with a promising development plan | 5. Working proof of concept

*Mark only one oval.*

| 1 | 2 | 3 | 4 | 5 |
|---|---|---|---|---|
| ◯ | ◯ | ◯ | ◯ | ◯ |

17. What impact would more funding have on the projects potential progress? *

1. No change in progress allowed by more funding | 3. Funding potentially could help, but other funding sources make JOGL funding obsolete | 5. Extra funding seems essential for the project to continue

*Mark only one oval.*

| 1 | 2 | 3 | 4 | 5 |
|---|---|---|---|---|
| ◯ | ◯ | ◯ | ◯ | ◯ |

#### Project Type

18. Is this project a product or data analysis/model *

*Mark only one oval.*

- ◯ A data analysis project/model *Skip to question 26*
- ◯ A Software/Hardware product/New Method/Research Eg: Diagnostic test
- ◯ Other: ____________________

#### Product Impact

19. Fit between the project's approach/methodology and the problem they have stated to resolve *

1. Inappropriate | 3. Appropriate | 5.Very appropriate

*Mark only one oval.*

| 1 | 2 | 3 | 4 | 5 |
|---|---|---|---|---|
| ◯ | ◯ | ◯ | ◯ | ◯ |

20. How relevant is the project to the COVID-19 crisis and alleviation of suffering? *

1. Not at all | 3. Moderately | 5. Essential

*Mark only one oval.*

| 1 | 2 | 3 | 4 | 5 |
|---|---|---|---|---|
| ◯ | ◯ | ◯ | ◯ | ◯ |

21. The scalability of the project in the long term, taking manufacturing into consideration *

1. No sustainability model and no scaling potential | 3. Sustainable and scalable but still unstructured plan | 5. Already applying a sustainable plan & good scalability

*Mark only one oval.*

| 1 | 2 | 3 | 4 | 5 |
|---|---|---|---|---|
| ⬭ | ⬭ | ⬭ | ⬭ | ⬭ |

22. How well has the project addressed concerns such as quality testing? *

1. Not well | 3. Clear but not entirely appropriate | 5. Very clear and relevant

*Mark only one oval.*

| 1 | 2 | 3 | 4 | 5 |
|---|---|---|---|---|
| ⬭ | ⬭ | ⬭ | ⬭ | ⬭ |

23. The project's dissemination strategy (quality of documentation for goals, results, methods and needs; open access model, outreach) is this/will this project be easily reproducible? *

1. Irreproducible and poorly documented 3. Good documentation but with few weaknesses 5. Easily reproducible & very well communicated

*Mark only one oval.*

| 1 | 2 | 3 | 4 | 5 |
|---|---|---|---|---|
| ⬭ | ⬭ | ⬭ | ⬭ | ⬭ |

24. Project Originality *

How much is this app/device needed? Are there already apps or devices that can do this job in an open source way? 1. The project is unoriginal and done before | 3. It has been done before, but not in an open source way | 5. The project is completely original as far as the reviewer is aware.

*Mark only one oval.*

| 1 | 2 | 3 | 4 | 5 |
|---|---|---|---|---|
| ⬭ | ⬭ | ⬭ | ⬭ | ⬭ |

25. Project practicality: Although the design quality may be high, there may be more suitable alternatives. Is this the case? *

1. Many more suitable alternatives exist | 3. The design or research has advantages to current designs that could be useful long term | 5. The design or research surpasses current research, or improves current efficiency

*Mark only one oval.*

| 1 | 2 | 3 | 4 | 5 |
|---|---|---|---|---|
| ⬭ | ⬭ | ⬭ | ⬭ | ⬭ |

*Skip to question 34*

#### Data Analysis project Impact

26. Fit between the project's approach/methodology and the problem they have stated to resolve *

1. Inappropriate | 3. Appropriate | 5.Very appropriate

*Mark only one oval.*

| 1 | 2 | 3 | 4 | 5 |
|---|---|---|---|---|
| ⬭ | ⬭ | ⬭ | ⬭ | ⬭ |

27. How relevant will the result of this project be for policy makers? *

1. Not at all | 3. Moderately | 5. Essential

*Mark only one oval.*

| 1 | 2 | 3 | 4 | 5 |
|---|---|---|---|---|
| ⬭ | ⬭ | ⬭ | ⬭ | ⬭ |

28. How relevant will the result of this project be for the public? *

1. Not at all | 3. Moderately | 5. Essential

*Mark only one oval.*

| 1 | 2 | 3 | 4 | 5 |
|---|---|---|---|---|
| ⬭ | ⬭ | ⬭ | ⬭ | ⬭ |

29. The project's dissemination strategy (quality of documentation for goals, results, methods and needs; open access model, outreach) how reproducible does this project seem? *

1. Irreproducible and poorly documented 3. Good documentation but with few weaknesses 5. Easily reproducible & very well communicated

*Mark only one oval.*

| 1 | 2 | 3 | 4 | 5 |
|---|---|---|---|---|
| ⬭ | ⬭ | ⬭ | ⬭ | ⬭ |

30. How accessible will the result of the project be to the general public? *

*Mark only one oval.*

| | 1 | 2 | 3 | 4 | 5 | |
|---|---|---|---|---|---|---|
| Unaccessible | ⬭ | ⬭ | ⬭ | ⬭ | ⬭ | Easily accessible and distributed on media |

31. How accessible will the result of the project be to policy makers? *

*Mark only one oval.*

| | 1 | 2 | 3 | 4 | 5 | |
|---|---|---|---|---|---|---|
| Unaccessible | ⬭ | ⬭ | ⬭ | ⬭ | ⬭ | Easily accessible and distributed to policy makers specifically |

32. Project Originality *

How much is this analysis needed/unique? Are there already open source analysis's pipelines out there that can do this? 1. Unoriginal and done before/not needed | 3.Done before but not relevant to this crisis | 5. The project is original as far as the reviewer is aware.

*Mark only one oval.*

| 1 | 2 | 3 | 4 | 5 |
|---|---|---|---|---|
| ◯ | ◯ | ◯ | ◯ | ◯ |

33. Project practicality: Although the design quality may be high, there may be more suitable alternatives. Is this the case? *

1. Many more suitable alternatives exist | 3. The design or research has advantages to current designs that could be useful long term | 5. The design or research surpasses current research, or improves current efficiency

*Mark only one oval.*

| 1 | 2 | 3 | 4 | 5 |
|---|---|---|---|---|
| ◯ | ◯ | ◯ | ◯ | ◯ |

#### Additional comments and reviewer self-evaluation

You may provide below additional comments addressed to the team leaders of the project, and private comments to the admin team.

Please, also assess your expertise in the fields relevant to evaluate the project.

34. Are they a previous applicant applying for further funding? (Indicated by presence of Section 8/ Progress since JOGL funds recieved) *

*Mark only one oval.*

◯ Yes

◯ No

35. [To the team leaders] Other general constructive comments about the project *

36. [To the team leaders] Other general critical comments about the project, that could be improved *

37. [To admins] Private comment - anything you would like to inform the admin team of, privately.

38. What is your expertise that is relevant to evaluate this project? *

Please provide justification of your expertise if possible.

39. How would you assess your own abilities to review this project? *

*Mark only one oval.*

| | 1 | 2 | 3 | 4 | 5 | |
|---|---|---|---|---|---|---|
| I cannot be trusted to assess this project, and did not fully grasp the project. | ⬭ | ⬭ | ⬭ | ⬭ | ⬭ | I can fully assess this project, based on my pı |

40. Are you part of the project team that you are currently assessing? *

*Mark only one oval.*

- Yes
- No

41. Are you part of a project team within the OpenCOVID19 initiative? *

*Mark only one oval.*

- Yes
- No

42. Would you like us to include you in our "Experts community" and provide similar reviews to projects when you have time in the future? *

*Mark only one oval.*

- Yes
- No

43. What is your JOGL username ?(if applicable)

44. What is your Slack Handle on the OpenCovid19 slack?(if applicable)

This content is neither created nor endorsed by Google.

Google Forms
