## Supplementary material for "Community review: a robust and scalable selection system for resource allocation within open science and innovation communities": Review form JOGL 5

### Project Technical Evaluation Form

Thank you for participating in our peer review process! Please find your selection of projects to review in an email from us if you are an applicant. For volunteer reviewers, here is a link to the projects for all reviewers page: https://tinyurl.com/ReviewJOGL. Take a look at all our projects and find the ones you would like to review most!

As an applicant due to the number of applicants, and to keep the process sustainable and truly open, you must review four other projects that you feel suit your skills most. This is up to the team leaders to complete. You cannot review your own project. Please fill your JOGL profile as much as possible, as this will let us improve the review process in future rounds by expertise matching.

As a volunteer reviewer, Thank you! Unaffiliated with a project you can use any email address, but to stop project bias you must declare any affiliations, cannot mark the work of your peers, must review at least three projects, and also create a JOGL account for your review to count. This is all to prevent misconduct in the process.

A new form is needed to review each project. The review process should take around 20 minutes or longer per project, but in some cases it might be remarkably quick. Only emails listed as project participants on JOGL will be counted in the review process, unless a specific email reviews at least 3 projects thoroughly and within expected bounds (as moderated by other review responses). This allows enthusiastic volunteer reviewers to take part in the process, but prevents malicious and bias reviewing.

You can find all the necessary information on each particular project proposal in the "About" section of their project page, or attached in their "Documents" section. This form is used for all quantitative measurements of projects therefore ultimately determines which projects receive a grant. In addition if you have time, to benefit project teams as greatly as possible, please leave a short positive note and a hello on the project's newsfeed (you may have to symbolically join their team as a reviewer to be able to post, but thats what open science is about). If this is not possible due to project permissions, use this review form for that feedback.

Feel free to connect with other projects to form collaborations, many projects share some of the same goals, and teams have joined forces in the past.Thank you for taking part in open science and deciding who will receive funding.

 with any issues or questions.

---

*Obligatoire

1. Adresse e-mail *

_______________________________

2. Reviewer JOGL profile URL *

For your review to count and prevent misconduct, you must provide your JOGL profile url Eg: https://app.jogl.io/user/3/Tholand. Create an account here if you have not already: https://app.jogl.io/signup. Click your profile picture/icon to access this page.

_______________________________

3. Reviewer Agreement *

We plan on using the anonymised reviewing behaviour in our rounds to form new ways to review. We also plan to publish this anonymised data in order to ensure the reviewing process is open. Do you consent to us analysing and reporting your anonymised expertise, and the reviewing behaviour displayed in this form?

*Une seule réponse possible.*

- [ ] Yes.

#### Code of Conduct

#### Code of Conduct

Read the article fully – please read the full text of the article and view all associated figures, tables and data;

Be thorough – a peer review regards the project in full as well as the individual points asked;

Be specific – your comments to other projects at the end, should contain as much detail as possible, with references where appropriate, so the authors are able to fully address the issue and improve for next round, or establish a better project;

Be constructive in your criticism – do not hesitate to include any concerns or criticisms you may have in your review, however, please do so in a constructive and respectful manner;

Avoid derogatory comments or tone – review as you wish to be reviewed and ensure that your comments focus on the scientific content of the article in question rather than the authors themselves.

4. URL of the project to be reviewed *

Copy and paste the URL of the project you are reviewing in this form.

5. Do you confirm you are not directly involved with the project you are about to review? *

Reminder: you cannot review your own project or a project that you have been involved with

*Une seule réponse possible.*

- No
- Yes

#### Project differentiation

6. Has this project been funded by JOGL before? *

Projects previously funded by JOGL will have a green badge adjacent to their name upon inspection indicating "This project has been reviewed well by a JOGL hosted Open Peer Review"

*Une seule réponse possible.*

- Yes
- No *Passer à la question 68*

#### Previous Reviewers

In order to save time, and ask less questions of you, we are asking if you have reviewed this project in previous rounds already

7. Have you reviewed this project before in previous rounds?

*Une seule réponse possible.*

- Yes
- No *Passer à la question 36*

#### Feasibility

This section is designed to assess the feasibility of already funded projects.

8. Project Timeline: How achievable are the new project objectives within 3 months? *

1. The new objectives are not achievable | 3. The new objectives are achievable, but will take a great effort by all involved and more participants | 5. The new objectives are very achievable with little extra input required.

*Une seule réponse possible.*

| 1 | 2 | 3 | 4 | 5 |
|---|---|---|---|---|
| ○ | ○ | ○ | ○ | ○ |

9. What is the quality of the budgeting for the project's new objectives? *

1. Low Quality/Non existent. 3. Costing exists and has specific costs allocated. 5. Costing is fully transparent and documented.

*Une seule réponse possible.*

| 1 | 2 | 3 | 4 | 5 |
|---|---|---|---|---|
| ○ | ○ | ○ | ○ | ○ |

10. Is the project's current team composition sufficient to achieve the new objectives set? *

1. Project team members inexperienced, or no details of team given | 3. The team expertise seem sufficient to achieve the goals set out by the project | 5. The project has multiple highly distinguished and or/experienced members who have a history of completing similar projects.

*Une seule réponse possible.*

| 1 | 2 | 3 | 4 | 5 |
|---|---|---|---|---|
| ○ | ○ | ○ | ○ | ○ |

11. Does the project have access to the materials/ facilities required to complete the new objectives set? *

1. The project does not have access to the materials or facilities required| 3. The project has access to materials/facilities, however not enough to fully scale the project | 5. The project has enough facility access to be completed to fruition.

*Une seule réponse possible.*

| 1 | 2 | 3 | 4 | 5 |
|---|---|---|---|---|
| ○ | ○ | ○ | ○ | ○ |

12. How clear is the project proposal, on its new objectives for this round? *

1. Unclear and/or irrelevant | 3. Clear, but relevant points are missing | 5. Very clear and relevant

*Une seule réponse possible.*

| 1 | 2 | 3 | 4 | 5 |
|---|---|---|---|---|
| ○ | ○ | ○ | ○ | ○ |

13. Project design I: What is the project design quality for their new objectives? Are there flaws? *

1. Many problems with project's new objectives in their design | 3. Design seems feasible, but it's inefficient | 5. Brilliant Design, well documented.

*Une seule réponse possible.*

| 1 | 2 | 3 | 4 | 5 |
|---|---|---|---|---|
| ⬭ | ⬭ | ⬭ | ⬭ | ⬭ |

14. Project design II: References and Prior Research, is there good foundation for this project? (Not applicable to some hardware of humanitarian projects)

1. No references or reference to published research or projects (if applicable) | 3. References poor in relevance, or their own preliminary data not shown (If applicable) | 5. Multiple relevant references, and/or preliminary data shown (If applicable)

*Une seule réponse possible.*

| 1 | 2 | 3 | 4 | 5 |
|---|---|---|---|---|
| ⬭ | ⬭ | ⬭ | ⬭ | ⬭ |

15. Which category would you say the project falls under? *

*Une seule réponse possible.*

- ⬭ Medical Hardware
- ⬭ Self Screening (Software)
- ⬭ Other Software/Appps
- ⬭ Diagnostic technology
- ⬭ Data Analysis/Research/Modelling project
- ⬭ Personal Protective equipment
- ⬭ Education
- ⬭ Autre : ______________________

16. Clarity and relevance of the project's new objectives, its timeline and it's needs for future (major tasks, milestones) *

1. Unclear and irrelevant | 3. Clear but relevant points are missing | 5. Very clear and relevant

*Une seule réponse possible.*

| 1 | 2 | 3 | 4 | 5 |
|---|---|---|---|---|
| ⬭ | ⬭ | ⬭ | ⬭ | ⬭ |

17. If this project is biological in nature, do they address the biosafety and regulations requirements? (if applicable)

1. Not at all | 3. Aware but unclear integration | 5. Actively engaged and addressed these regulations

*Une seule réponse possible.*

| 1 | 2 | 3 | 4 | 5 |
|---|---|---|---|---|
| ⬭ | ⬭ | ⬭ | ⬭ | ⬭ |

18. If it is a data driven project how feasible is the data collection for this project ? (Really Hard / The data exists already) (Data analytics projects only)

1. Not at all | 3. Aware but unclear integration | 5. Actively engaged and addressed these regulations

*Une seule réponse possible.*

| 1 | 2 | 3 | 4 | 5 |
|---|---|---|---|---|
| ⬭ | ⬭ | ⬭ | ⬭ | ⬭ |

19. What is the project's state of progress? *

1. No results yet | 3. Some results but with a promising development plan | 5. Working proof of concept

*Une seule réponse possible.*

| 1 | 2 | 3 | 4 | 5 |
|---|---|---|---|---|
| ⬭ | ⬭ | ⬭ | ⬭ | ⬭ |

20. What impact would more funding have on the projects potential progress? *

1. No change in progress allowed by more funding | 3. Funding potentially could help, but other funding sources make JOGL funding obsolete | 5. Extra funding seems essential for the project to continue

*Une seule réponse possible.*

| 1 | 2 | 3 | 4 | 5 |
|---|---|---|---|---|
| ⬭ | ⬭ | ⬭ | ⬭ | ⬭ |

#### Project Type

21. Is this project a product/research project or data analysis/model of existing data *

*Une seule réponse possible.*

- ⬭ A data analysis project/model *Passer à la question 28*
- ⬭ A Software/Hardware product/New Method/Research Eg: Diagnostic test
- ⬭ Autre : ______________________

#### Product Impact

This section is designed to assess the impact of already funded projects.

22. Fit between the project's new objectives and the problem they have stated to resolve *

1. Inappropriate | 3. Appropriate | 5.Very appropriate

*Une seule réponse possible.*

| 1 | 2 | 3 | 4 | 5 |
|---|---|---|---|---|
| ⬭ | ⬭ | ⬭ | ⬭ | ⬭ |

23. How relevant is the projects new objectives to the COVID-19 crisis and alleviation of suffering? *

1. Not at all | 3. Moderately | 5. Essential

*Une seule réponse possible.*

| 1 | 2 | 3 | 4 | 5 |
|---|---|---|---|---|
| ⬭ | ⬭ | ⬭ | ⬭ | ⬭ |

24. The scalability of the project in the long term, taking manufacturing into consideration *

1. No sustainability model and no scaling potential | 3. Sustainable and scalable, but still unstructured plan | 5. The project is already applying a sustainable plan & has good scalability

*Une seule réponse possible.*

| 1 | 2 | 3 | 4 | 5 |
|---|---|---|---|---|
| ⬭ | ⬭ | ⬭ | ⬭ | ⬭ |

25. How well has the project addressed concerns such as quality testing? (If applicable) Especially applicable to PPE.

1. Not well/no mention of quality testing, despite this being essential | 3. Quality testing is clear, but not entirely appropriate | 5. Quality testing is very clear and relevant.

*Une seule réponse possible.*

| 1 | 2 | 3 | 4 | 5 |
|---|---|---|---|---|
| ⬭ | ⬭ | ⬭ | ⬭ | ⬭ |

26. Has the project been openly sharing its results so far (its documentation for goals, results, methods and needs; open access model, outreach) is this/will this project be easily reproducible? *

1. It is irreproducible and poorly documented, or has shared no results 3. The results have good documentation, but with few weaknesses 5. This project results are open, easily reproducible [if applicable] & very well communicated

*Une seule réponse possible.*

| 1 | 2 | 3 | 4 | 5 |
|---|---|---|---|---|
| ⬭ | ⬭ | ⬭ | ⬭ | ⬭ |

27. Project practicality: Although the design quality may be high, there may be more suitable alternatives. Is this the case? *

1. Many more suitable alternatives exist | 3. The design or research has advantages to current designs that could be useful long term | 5. The design or research surpasses current research, or improves current efficiency

*Une seule réponse possible.*

| 1 | 2 | 3 | 4 | 5 |
|---|---|---|---|---|
| ⬭ | ⬭ | ⬭ | ⬭ | ⬭ |

*Passer à la question 66*

Data Analysis project Impact

28. Fit between the project's new objectives and the problem they have stated to resolve *

1. Inappropriate | 3. Appropriate | 5.Very appropriate

*Une seule réponse possible.*

| 1 | 2 | 3 | 4 | 5 |
|---|---|---|---|---|
| ◯ | ◯ | ◯ | ◯ | ◯ |

29. How relevant will the new results of this project be for policy makers? *

1. Not at all | 3. Moderately | 5. Essential

*Une seule réponse possible.*

| 1 | 2 | 3 | 4 | 5 |
|---|---|---|---|---|
| ◯ | ◯ | ◯ | ◯ | ◯ |

30. How relevant will the new results of this project be for the public? *

1. Not at all | 3. Moderately | 5. Essential

*Une seule réponse possible.*

| 1 | 2 | 3 | 4 | 5 |
|---|---|---|---|---|
| ◯ | ◯ | ◯ | ◯ | ◯ |

31. The project's dissemination strategy (quality of documentation for goals, results, methods and needs; open access model, outreach) how reproducible is this project? *

1. The project is currently Irreproducible and poorly documented 3. The project has shared results, has good documentation but with few weaknesses 5. The project is easily reproducible & very well communicated

*Une seule réponse possible.*

| 1 | 2 | 3 | 4 | 5 |
|---|---|---|---|---|
| ◯ | ◯ | ◯ | ◯ | ◯ |

32. How accessible is the results of the project so far to the general public? *

*Une seule réponse possible.*

| | 1 | 2 | 3 | 4 | 5 | |
|---|---|---|---|---|---|---|
| Unaccessible | ◯ | ◯ | ◯ | ◯ | ◯ | Easily accessible and distributed on media |

33. How accessible are the results of the project be to policy makers? *

*Une seule réponse possible.*

| | 1 | 2 | 3 | 4 | 5 | |
|---|---|---|---|---|---|---|
| Unaccessible | ◯ | ◯ | ◯ | ◯ | ◯ | Easily accessible and distributed to policy makers specifically |

34. Project Originality *

How much is this new analysis needed/unique? Are there already open source analysis's pipelines out there that can do this? 1. Unoriginal and done before/not needed | 3.Done before but not relevant to this crisis | 5. The project is original as far as the reviewer is aware.

*Une seule réponse possible.*

| 1 | 2 | 3 | 4 | 5 |
|---|---|---|---|---|
| ◯ | ◯ | ◯ | ◯ | ◯ |

35. Project practicality: Although the design quality may be high, there may be more suitable alternatives. Is this the case? *

1. Many more suitable alternatives exist | 3. The design or research has advantages to current designs that could be useful long term | 5. The design or research surpasses current research, or improves current efficiency

*Une seule réponse possible.*

| 1 | 2 | 3 | 4 | 5 |
|---|---|---|---|---|
| ◯ | ◯ | ◯ | ◯ | ◯ |

*Passer à la question 66*

#### Feasibility

This section is designed to assess the feasibility of already funded projects.

36. Project Timeline: How achievable are the new project objectives within 3 months? *

1. The new objectives are not achievable | 3. The new objectives are achievable, but will take a great effort by all involved and more participants | 5. The new objectives are very achievable with little extra input required.

*Une seule réponse possible.*

| 1 | 2 | 3 | 4 | 5 |
|---|---|---|---|---|
| ◯ | ◯ | ◯ | ◯ | ◯ |

37. What is the quality of the budgeting for the project's new objectives? *

1. Low Quality/Non existent. 3. Costing exists and has specific costs allocated. 5. Costing is fully transparent and documented.

*Une seule réponse possible.*

| 1 | 2 | 3 | 4 | 5 |
|---|---|---|---|---|
| ◯ | ◯ | ◯ | ◯ | ◯ |

38. Is the project's current team composition sufficient to achieve the new objectives set? *

1. Project team members inexperienced, or no details of team given | 3. The team expertise seem sufficient to achieve the goals set out by the project | 5. The project has multiple highly distinguished and or/experienced members who have a history of completing similar projects.

*Une seule réponse possible.*

| 1 | 2 | 3 | 4 | 5 |
|---|---|---|---|---|
| ◯ | ◯ | ◯ | ◯ | ◯ |

39. Does the project have access to the materials/ facilities required to complete the new objectives set? *

1. The project does not have access to the materials or facilities required| 3. The project has access to materials/facilities, however not enough to fully scale the project | 5. The project has enough facility access to be completed to fruition.

*Une seule réponse possible.*

| 1 | 2 | 3 | 4 | 5 |
|---|---|---|---|---|
| ⬭ | ⬭ | ⬭ | ⬭ | ⬭ |

40. How clear is the project proposal, on its new objectives for this round? *

1. Unclear and/or irrelevant | 3. Clear, but relevant points are missing | 5. Very clear and relevant

*Une seule réponse possible.*

| 1 | 2 | 3 | 4 | 5 |
|---|---|---|---|---|
| ⬭ | ⬭ | ⬭ | ⬭ | ⬭ |

41. Project design I: What is the project design quality for their new objectives? Are there flaws? *

1. Many problems with project's new objectives in their design | 3. Design seems feasible, but it's inefficient | 5. Brilliant Design, well documented.

*Une seule réponse possible.*

| 1 | 2 | 3 | 4 | 5 |
|---|---|---|---|---|
| ⬭ | ⬭ | ⬭ | ⬭ | ⬭ |

42. Project design II: References and Prior Research, is there good foundation for this project? (Not applicable to some hardware of humanitarian projects)

1. No references or reference to published research or projects (if applicable) | 3. References poor in relevance, or their own preliminary data not shown (If applicable) | 5. Multiple relevant references, and/or preliminary data shown (If applicable)

*Une seule réponse possible.*

| 1 | 2 | 3 | 4 | 5 |
|---|---|---|---|---|
| ⬭ | ⬭ | ⬭ | ⬭ | ⬭ |

43. Which category would you say the project falls under? *

*Une seule réponse possible.*

- ⬭ Medical Hardware
- ⬭ Self Screening (Software)
- ⬭ Other Software/Appps
- ⬭ Diagnostic technology
- ⬭ Data Analysis/Research/Modelling project
- ⬭ Personal Protective equipment
- ⬭ Education
- ⬭ Autre : ____________________

44. Clarity and relevance of the project's new objectives, its timeline and it's needs for future (major tasks, milestones) *

1. Unclear and irrelevant | 3. Clear but relevant points are missing | 5. Very clear and relevant

*Une seule réponse possible.*

| 1 | 2 | 3 | 4 | 5 |
|---|---|---|---|---|
| ◯ | ◯ | ◯ | ◯ | ◯ |

45. If this project is biological in nature, do they address the biosafety and regulations requirements? (if applicable)

1. Not at all | 3. Aware but unclear integration | 5. Actively engaged and addressed these regulations

*Une seule réponse possible.*

| 1 | 2 | 3 | 4 | 5 |
|---|---|---|---|---|
| ◯ | ◯ | ◯ | ◯ | ◯ |

46. If it is a data driven project how feasible is the data collection for this project ? (Really Hard / The data exists already) (Data analytics projects only)

1. Not at all | 3. Aware but unclear integration | 5. Actively engaged and addressed these regulations

*Une seule réponse possible.*

| 1 | 2 | 3 | 4 | 5 |
|---|---|---|---|---|
| ◯ | ◯ | ◯ | ◯ | ◯ |

47. What is the project's state of progress? *

1. No results yet | 3. Some results but with a promising development plan | 5. Working proof of concept

*Une seule réponse possible.*

| 1 | 2 | 3 | 4 | 5 |
|---|---|---|---|---|
| ◯ | ◯ | ◯ | ◯ | ◯ |

48. What impact would more funding have on the projects potential progress? *

1. No change in progress allowed by more funding | 3. Funding potentially could help, but other funding sources make JOGL funding obsolete | 5. Extra funding seems essential for the project to continue

*Une seule réponse possible.*

| 1 | 2 | 3 | 4 | 5 |
|---|---|---|---|---|
| ◯ | ◯ | ◯ | ◯ | ◯ |

Project Type

49. Is this project a product/research project or data analysis/model of existing data *

*Une seule réponse possible.*

- A data analysis project/model *Passer à la question 58*
- A Software/Hardware product/New Method/Research Eg: Diagnostic test
- Autre : ______

#### Product Impact

This section is designed to assess the impact of already funded projects.

50. Fit between the original project and the problem they have stated to resolve *

1. Inappropriate | 3. Appropriate | 5.Very appropriate

*Une seule réponse possible.*

| 1 | 2 | 3 | 4 | 5 |
|---|---|---|---|---|

51. Fit between the project's new objectives and the problem they have stated to resolve *

1. Inappropriate | 3. Appropriate | 5.Very appropriate

*Une seule réponse possible.*

| 1 | 2 | 3 | 4 | 5 |
|---|---|---|---|---|

52. How relevant is the projects new objectives to the COVID-19 crisis and alleviation of suffering? *

1. Not at all | 3. Moderately | 5. Essential

*Une seule réponse possible.*

| 1 | 2 | 3 | 4 | 5 |
|---|---|---|---|---|

53. The scalability of the project in the long term, taking manufacturing into consideration *

1. No sustainability model and no scaling potential | 3. Sustainable and scalable, but still unstructured plan | 5. The project is already applying a sustainable plan & has good scalability

*Une seule réponse possible.*

| 1 | 2 | 3 | 4 | 5 |
|---|---|---|---|---|

54. How well has the project addressed concerns such as quality testing? (If applicable) Especially applicable to PPE.

1. Not well/no mention of quality testing, despite this being essential | 3. Quality testing is clear, but not entirely appropriate | 5. Quality testing is very clear and relevant.

*Une seule réponse possible.*

| 1 | 2 | 3 | 4 | 5 |
|---|---|---|---|---|
| ◯ | ◯ | ◯ | ◯ | ◯ |

55. Has the project been openly sharing its results so far (its documentation for goals, results, methods and needs; open access model, outreach) is this/will this project be easily reproducible? *

1. It is irreproducible and poorly documented, or has shared no results 3. The results have good documentation, but with few weaknesses 5. This project results are open, easily reproducible [if applicable] & very well communicated

*Une seule réponse possible.*

| 1 | 2 | 3 | 4 | 5 |
|---|---|---|---|---|
| ◯ | ◯ | ◯ | ◯ | ◯ |

56. Project Originality *

How much is this new analysis needed/unique? Are there already open source analysis's pipelines out there that can do this? 1. Unoriginal and done before/not needed | 3.Done before but not relevant to this crisis | 5. The project is original as far as the reviewer is aware.

*Une seule réponse possible.*

| 1 | 2 | 3 | 4 | 5 |
|---|---|---|---|---|
| ◯ | ◯ | ◯ | ◯ | ◯ |

57. Project practicality: Although the design quality may be high, there may be more suitable alternatives. Is this the case? *

1. Many more suitable alternatives exist | 3. The design or research has advantages to current designs that could be useful long term | 5. The design or research surpasses current research, or improves current efficiency

*Une seule réponse possible.*

| 1 | 2 | 3 | 4 | 5 |
|---|---|---|---|---|
| ◯ | ◯ | ◯ | ◯ | ◯ |

*Passer à la question 66*

#### Data Analysis project Impact

This investigates the impact of projects that have received JOGL funding.

58. Fit between the project's new objectives and the problem they have stated to resolve *

1. Inappropriate | 3. Appropriate | 5.Very appropriate

*Une seule réponse possible.*

| 1 | 2 | 3 | 4 | 5 |
|---|---|---|---|---|
| ◯ | ◯ | ◯ | ◯ | ◯ |

59. How relevant will the new results of this project be for policy makers? *

1. Not at all | 3. Moderately | 5. Essential

*Une seule réponse possible.*

| 1 | 2 | 3 | 4 | 5 |
|---|---|---|---|---|
| ⬭ | ⬭ | ⬭ | ⬭ | ⬭ |

60. How relevant will the new results of this project be for the public? *

1. Not at all | 3. Moderately | 5. Essential

*Une seule réponse possible.*

| 1 | 2 | 3 | 4 | 5 |
|---|---|---|---|---|
| ⬭ | ⬭ | ⬭ | ⬭ | ⬭ |

61. The project's dissemination strategy (quality of documentation for goals, results, methods and needs; open access model, outreach) how reproducible is this project? *

1. The project is currently Irreproducible and poorly documented 3. The project has shared results, has good documentation but with few weaknesses 5. The project is easily reproducible & very well communicated

*Une seule réponse possible.*

| 1 | 2 | 3 | 4 | 5 |
|---|---|---|---|---|
| ⬭ | ⬭ | ⬭ | ⬭ | ⬭ |

62. How accessible is the results of the project so far to the general public? *

*Une seule réponse possible.*

| | 1 | 2 | 3 | 4 | 5 | |
|---|---|---|---|---|---|---|
| Unaccessible | ⬭ | ⬭ | ⬭ | ⬭ | ⬭ | Easily accessible and distributed on media |

63. How accessible are the results of the project be to policy makers? *

*Une seule réponse possible.*

| | 1 | 2 | 3 | 4 | 5 | |
|---|---|---|---|---|---|---|
| Unaccessible | ⬭ | ⬭ | ⬭ | ⬭ | ⬭ | Easily accessible and distributed to policy makers specifically |

64. Project Originality *

How much is this new analysis needed/unique? Are there already open source analysis's pipelines out there that can do this? 1. Unoriginal and done before/not needed | 3.Done before but not relevant to this crisis | 5. The project is original as far as the reviewer is aware.

*Une seule réponse possible.*

| 1 | 2 | 3 | 4 | 5 |
|---|---|---|---|---|
| ⬭ | ⬭ | ⬭ | ⬭ | ⬭ |

65. Project practicality: Although the design quality may be high, there may be more suitable alternatives. Is this the case? *

1. Many more suitable alternatives exist | 3. The design or research has advantages to current designs that could be useful long term | 5. The design or research surpasses current research, or improves current efficiency

*Une seule réponse possible.*

| 1 | 2 | 3 | 4 | 5 |
|---|---|---|---|---|
| ◯ | ◯ | ◯ | ◯ | ◯ |

#### Section 8: Progress of previously funded projects

Impact assessment of previously funded projects, progress is tracked by Section 8.

66. How much has been achieved since the project was funded? *

1. The project hasn't made any progress clear/ has had no progress since being funded. 2. Little progress since last application/poor dissemination of progress. 3. Progress has been made and made clear by the proposal. 4. Great advances in the project such as websites, apps, pre-prints/papers and results are shown. 5. Progress is great, with multiple types of achievements made clear in the proposal's section 8.

*Une seule réponse possible.*

| 1 | 2 | 3 | 4 | 5 |
|---|---|---|---|---|
| ◯ | ◯ | ◯ | ◯ | ◯ |

67. Impact from project in relation to COVID19 *

1. No-one has benefitted from this project, and project has not progressed or impacted others 2. The cultural/academic impact from this project through or academic papers/media has likely had an impact and the project has progressed. 3. The project has achieved its goals, with cultural/scientific impact shown through participation in other events, small scale efforts to share the project or academic papers. 4. This project has already been used in the field, has participated in collaborative research papers and has been shared or used on a small scale to impact lives 5. This project has been used or has concrete plans to be used on a medium/large scale with impact so far in other aspects clearly shown throughout the proposal.

*Une seule réponse possible.*

| 1 | 2 | 3 | 4 | 5 |
|---|---|---|---|---|
| ◯ | ◯ | ◯ | ◯ | ◯ |

*Passer à la question 97*

#### Feasibility

This investigates the feasibility of projects yet to have received JOGL funding.

68. Project Timeline: How achievable is this project within 3 months? *

1. The project is not achievable | 3. The project is achievable, but will take a great effort by all involved and more participants | 5. The project is very achievable with little extra input required.

*Une seule réponse possible.*

| 1 | 2 | 3 | 4 | 5 |
|---|---|---|---|---|
| ◯ | ◯ | ◯ | ◯ | ◯ |

69. What is the quality of the project budgeting? *

1. Low Quality/Non existent. 3. Costing exists and has specific costs allocated. 5. Costing is fully transparent and documented.

*Une seule réponse possible.*

| 1 | 2 | 3 | 4 | 5 |
|---|---|---|---|---|
| ◯ | ◯ | ◯ | ◯ | ◯ |

70. Is the project's current team composition sufficient to achieve the projects goal? *

1. Project team members inexperienced, or no details of team given | 3. The team expertise seem sufficient to achieve the goals set out by the project | 5. The project has multiple highly distinguished and or/experienced members who have a history of completing similar projects.

*Une seule réponse possible.*

| 1 | 2 | 3 | 4 | 5 |
|---|---|---|---|---|
| ◯ | ◯ | ◯ | ◯ | ◯ |

71. Does the project have access to the materials/ facilities required to complete the project?

1. The project does not have access to the materials or facilities required| 3. The project has access to materials/facilities, however not enough to fully scale the project | 5. The project has enough facility access to be completed to fruition.

*Une seule réponse possible.*

| 1 | 2 | 3 | 4 | 5 |
|---|---|---|---|---|
| ◯ | ◯ | ◯ | ◯ | ◯ |

72. How clear is the project proposal? *

1. Unclear and/or irrelevant | 3. Clear, but relevant points are missing | 5. Very clear and relevant

*Une seule réponse possible.*

| 1 | 2 | 3 | 4 | 5 |
|---|---|---|---|---|
| ◯ | ◯ | ◯ | ◯ | ◯ |

73. Project design I: What is the project design quality? Are there flaws? *

1. Many problems with project design | 3. Design seems feasible, but it's inefficient | 5. Brilliant Design, well documented.

*Une seule réponse possible.*

| 1 | 2 | 3 | 4 | 5 |
|---|---|---|---|---|
| ◯ | ◯ | ◯ | ◯ | ◯ |

74. Project design II: References and Prior Research, is there good foundation for this project? (Not applicable to some hardware of humanitarian projects)

1. No references or reference to published research or projects (if applicable) | 3. References poor in relevance, or their own preliminary data not shown (If applicable) | 5. Multiple relevant references, and/or preliminary data shown.(If applicable)

*Une seule réponse possible.*

| 1 | 2 | 3 | 4 | 5 |
|---|---|---|---|---|
| ⬭ | ⬭ | ⬭ | ⬭ | ⬭ |

75. Which category would you say the project falls under? *

*Une seule réponse possible.*

- ⬭ Medical Hardware
- ⬭ Self Screening (Software)
- ⬭ Other Software/Appps
- ⬭ Diagnostic technology
- ⬭ Data Analysis/Research/Modelling project
- ⬭ Personal Protective equipment
- ⬭ Education
- ⬭ Autre : ____________________

76. Clarity and relevance of the project timeline and it's needs for future (major tasks, milestones) *

1. Unclear and irrelevant | 3. Clear but relevant points are missing | 5. Very clear and relevant

*Une seule réponse possible.*

| 1 | 2 | 3 | 4 | 5 |
|---|---|---|---|---|
| ⬭ | ⬭ | ⬭ | ⬭ | ⬭ |

77. If this project is biological in nature, do they address the biosafety and regulations requirements to address? (if applicable)

1. Not at all | 3. Aware but unclear integration | 5. Actively engaged and addressed these regulations

*Une seule réponse possible.*

| 1 | 2 | 3 | 4 | 5 |
|---|---|---|---|---|
| ⬭ | ⬭ | ⬭ | ⬭ | ⬭ |

78. If it is a data driven project how feasible is the data collection for this project ? (Really Hard / The data exists already) (Data analytics projects only)

1. Not at all | 3. Aware but unclear integration | 5. Actively engaged and addressed these regulations

*Une seule réponse possible.*

| 1 | 2 | 3 | 4 | 5 |
|---|---|---|---|---|
| ⬭ | ⬭ | ⬭ | ⬭ | ⬭ |

79. What is the project's state of progress? *

1. No results yet | 3. Some results but with a promising development plan | 5. Working proof of concept

*Une seule réponse possible.*

| | 1 | 2 | 3 | 4 | 5 |
|---|---|---|---|---|---|
| | ◯ | ◯ | ◯ | ◯ | ◯ |

80. What impact would more funding have on the projects potential progress? *

1. No change in progress allowed by more funding | 3. Funding potentially could help, but other funding sources make JOGL funding obsolete | 5. Extra funding seems essential for the project to continue

*Une seule réponse possible.*

| | 1 | 2 | 3 | 4 | 5 |
|---|---|---|---|---|---|
| | ◯ | ◯ | ◯ | ◯ | ◯ |

*Passer à la question 81*

#### Project Type

81. Is this project a product or data analysis/model *

*Une seule réponse possible.*

- ◯ A data analysis project/model using existing data *Passer à la question 89*
- ◯ A Software/Hardware product/New Method/Research Eg: Diagnostic test
- ◯ Autre : ____________________

*Passer à la question 81*

#### Product Impact

This investigates the impact of projects yet to have received JOGL funding.

82. Fit between the project's approach/methodology and the problem they have stated to resolve *

1. Inappropriate | 3. Appropriate | 5.Very appropriate

*Une seule réponse possible.*

| | 1 | 2 | 3 | 4 | 5 |
|---|---|---|---|---|---|
| | ◯ | ◯ | ◯ | ◯ | ◯ |

83. How relevant is the project to the COVID-19 crisis and alleviation of suffering? *

1. Not at all | 3. Moderately | 5. Essential

*Une seule réponse possible.*

| | 1 | 2 | 3 | 4 | 5 |
|---|---|---|---|---|---|
| | ◯ | ◯ | ◯ | ◯ | ◯ |

84. The scalability of the project in the long term, taking manufacturing into consideration *

1. There is no sustainability model and no scaling potential | 3. It is sustainable and scalable, but there is an unstructured plan | 5. The project is already applying a sustainable plan & has good scalability

*Une seule réponse possible.*

| 1 | 2 | 3 | 4 | 5 |
|---|---|---|---|---|
| ⬭ | ⬭ | ⬭ | ⬭ | ⬭ |

85. How well has the project addressed concerns such as quality testing? (If relevant) Especially relevant to diagnostic and PPE projects.

1. Not well/no quality testing | 3. Clear, but not entirely appropriate | 5. Very clear and relevant techniques on quality testing are used.

*Une seule réponse possible.*

| 1 | 2 | 3 | 4 | 5 |
|---|---|---|---|---|
| ⬭ | ⬭ | ⬭ | ⬭ | ⬭ |

86. The project's dissemination strategy (quality of documentation for goals, results, methods and needs; open access model, outreach) is this/will this project be easily reproducible? *

1. Irreproducible and poorly documented 3. Good documentation but with few weaknesses 5. Easily reproducible & very well communicated

*Une seule réponse possible.*

| 1 | 2 | 3 | 4 | 5 |
|---|---|---|---|---|
| ⬭ | ⬭ | ⬭ | ⬭ | ⬭ |

87. Project Originality *

How much is this app/device needed? Are there already apps or devices that can do this job in an open source way? 1. The project is unoriginal and done before | 3. It has been done before, but not in an open source way | 5. The project is completely original as far as the reviewer is aware.

*Une seule réponse possible.*

| 1 | 2 | 3 | 4 | 5 |
|---|---|---|---|---|
| ⬭ | ⬭ | ⬭ | ⬭ | ⬭ |

88. Project practicality: Although the design quality may be high, there may be more suitable alternatives. Is this the case? *

1. Many more suitable alternatives exist | 3. The design or research has advantages to current designs that could be useful long term | 5. The design or research surpasses current research, or improves current efficiency

*Une seule réponse possible.*

| 1 | 2 | 3 | 4 | 5 |
|---|---|---|---|---|
| ⬭ | ⬭ | ⬭ | ⬭ | ⬭ |

*Passer à la question 97*

#### Data Analysis project Impact

This investigates the impact of projects yet to have received JOGL funding.

89. Fit between the project's approach/methodology and the problem they have stated to resolve *

1. Inappropriate | 3. Appropriate | 5.Very appropriate

*Une seule réponse possible.*

| 1 | 2 | 3 | 4 | 5 |
|---|---|---|---|---|
| ◯ | ◯ | ◯ | ◯ | ◯ |

90. How relevant will the result of this project be for policy makers? *

1. Not at all | 3. Moderately | 5. Essential

*Une seule réponse possible.*

| 1 | 2 | 3 | 4 | 5 |
|---|---|---|---|---|
| ◯ | ◯ | ◯ | ◯ | ◯ |

91. How relevant will the result of this project be for the public? *

1. Not at all | 3. Moderately | 5. Essential

*Une seule réponse possible.*

| 1 | 2 | 3 | 4 | 5 |
|---|---|---|---|---|
| ◯ | ◯ | ◯ | ◯ | ◯ |

92. The project's dissemination strategy (quality of documentation for goals, results, methods and needs; open access model, outreach) how reproducible does this project seem? *

1. Irreproducible and poorly documented 3. Good documentation but with few weaknesses 5. Easily reproducible & very well communicated

*Une seule réponse possible.*

| 1 | 2 | 3 | 4 | 5 |
|---|---|---|---|---|
| ◯ | ◯ | ◯ | ◯ | ◯ |

93. How accessible will the result of the project be to the general public? *

*Une seule réponse possible.*

| | 1 | 2 | 3 | 4 | 5 | |
|---|---|---|---|---|---|---|
| Unaccessible | ◯ | ◯ | ◯ | ◯ | ◯ | Easily accessible and distributed on media |

94. How accessible will the result of the project be to policy makers? *

*Une seule réponse possible.*

| | 1 | 2 | 3 | 4 | 5 | |
|---|---|---|---|---|---|---|
| Unaccessible | ◯ | ◯ | ◯ | ◯ | ◯ | Easily accessible and distributed to policy makers specifically |

95. Project Originality *

How much is this analysis needed/unique? Are there already open source analysis's pipelines out there that can do this? 1. Unoriginal and done before/not needed | 3.Done before but not relevant to this crisis | 5. The project is original as far as the reviewer is aware.

*Une seule réponse possible.*

| 1 | 2 | 3 | 4 | 5 |
|---|---|---|---|---|
| ◯ | ◯ | ◯ | ◯ | ◯ |

96. Project practicality: Although the design quality may be high, there may be more suitable alternatives. Is this the case? *

1. Many more suitable alternatives exist | 3. The design or research has advantages to current designs that could be useful long term | 5. The design or research surpasses current research, or improves current efficiency

*Une seule réponse possible.*

| 1 | 2 | 3 | 4 | 5 |
|---|---|---|---|---|
| ◯ | ◯ | ◯ | ◯ | ◯ |

#### Additional comments and reviewer self-evaluation

You may provide below additional comments addressed to the team leaders of the project, and private comments to the admin team.

Please, also assess your expertise in the fields relevant to evaluate the project.

97. Overall Impression

What would you give the project proposal, in the context of relevance to COVID19 relief/research, it's quality and impact as an overall unspecific score? 1.An unremarkable project/irrelevant 2.Problems with quality/impact and relevance. 3. A good project, but lacking in areas of thought/quality 4. An impactful project with high quality proposal and relevance 5. A truly extraordinary project in terms of impact, feasibility and quality.

*Une seule réponse possible.*

| 1 | 2 | 3 | 4 | 5 |
|---|---|---|---|---|
| ◯ | ◯ | ◯ | ◯ | ◯ |

98. [To the team leaders privately] General critical comments about the project, that could be improved *

99. [To admins] Private comment - anything you would like to inform the admin team of privately that may concern JOGL.

100. What is your expertise that is relevant to evaluate this project? *

Please provide justification of your expertise if possible.

101. How would you assess your own abilities to review this project? *

*Une seule réponse possible.*

| | 1 | 2 | 3 | 4 | 5 | |
|---|---|---|---|---|---|---|
| I cannot be trusted to assess this project, and did not fully grasp the project. | | | | | | I can fully assess this project, based o[illegible] |

102. Are you part of a project team being reviewed this grant round? *

*Une seule réponse possible.*

- Yes
- No

103. Have you ever been part of a project team within the OpenCOVID19 initiative? *

*Une seule réponse possible.*

- Yes
- No

104. Conflict of interest

Is there any non-academic reason you might have marked this project differently to other reviewers? What is that reason? We analyse reviewing behaviour and these conflicts are important for the evaluation of this pilot scheme.

105. Would you like us to include you in our "Experts community" and provide similar reviews to projects when you have time in the future?

*Une seule réponse possible.*

- Yes
- No

#### Written Feedback for projects

Finally, we encourage reviewers to directly comment any positive aspects of the projects they assess as well as constructive help on the project's newsfeed channel. Thank you for taking part in the review, and we hope you find the comments other users provide your project useful, if you are an applicant.

Good luck and thank you for taking part in the beginnings of truly open science.

Ce contenu n'est ni rédigé, ni cautionné par Google.

Google Forms
