## Supplementary material for "Community review: a robust and scalable selection system for resource allocation within open science and innovation communities": Helpful Engineering Review Form

### Project Technical Evaluation Form

Thank you for participating in our peer review process! Find the project list to review here https://docs.google.com/spreadsheets/d/1A3-GXy6gekkMAw4EtaC6QkmvWcNPPPuwUJdNkvmb1d4/edit?usp=sharing Review as many projects as you can, but don't feel as though you must review them all.

* Required

1. Email *

_______________

2. Project number *

You can find the list of links and names of projects to review here: https://docs.google.com/spreadsheets/d/1A3-GXy6gekkMAw4EtaC6QkmvWcNPPPuwUJdNkvmb1d4/edit?usp=sharing

_______________

#### Feasibility

3. Development Timeframe: Reviewer estimate : Leave blank it unsure

*Mark only one oval.*

- ◯ less than a week
- ◯ less than 6 Weeks
- ◯ less than 3 months
- ◯ less than 6 months
- ◯ Greater than 6 months

4. Project Timeline: How achievable is this project within 6 months? *

1. The project is not achievable 3. The project is achievable, but will take a great effort by all involved and more participants. 5. The project is very achievable with little input required.

*Mark only one oval.*

| 1 | 2 | 3 | 4 | 5 |
|---|---|---|---|---|
| ◯ | ◯ | ◯ | ◯ | ◯ |

5. Build cost estimate per unit (if applicable)

*Mark only one oval.*

- ◯ less than 100
- ◯ less than 500
- ◯ less than 1000
- ◯ less than 5000

6. Build cost currency (if applicable)

*Mark only one oval.*

- ◯ USD
- ◯ Euro

#### Approachability

What tools and skills are need to build this project?

7. Build skill difficulty *

*Mark only one oval.*

- ( ) Crafting
- ( ) Hand power tools
- ( ) Digital tools
- ( ) Manufacturer
- ( ) Medical knowledge

8. Material Sources (if applicable)

*Check all that apply.*

- [ ] Home
- [ ] Retail stores
- [ ] Online Retail
- [ ] Industrial
- [ ] Medical grade

#### Project Approach

In this section, you will assess the approach of the project. Please rank from 1 to 5 each criteria below. You can provide additional comments in the appropriate field.

9. The clarity of the project proposal *

1. Unclear and/or irrelevant | 3. Clear but relevant points are missing | 5. Very clear and relevant

*Mark only one oval.*

| 1 | 2 | 3 | 4 | 5 |
|---|---|---|---|---|
| ( ) | ( ) | ( ) | ( ) | ( ) |

10. The clarity of the project proposal [Comment, optional]

Please, comment and explain the score you have given. You can also give constructive feedback to help the project improve on this criteria.

11. Project Design: What is the project design quality? are there flaws? [Score] *

1. Many problems with project design. 3. Design seems feasible but its inefficient 5. Brilliant Design, well documented.

*Mark only one oval.*

| 1 | 2 | 3 | 4 | 5 |
|---|---|---|---|---|
| ⬭ | ⬭ | ⬭ | ⬭ | ⬭ |

12. Project Design: What is the project design quality? are there flaws [Comments, optional]

13. Fit between the project's approach/methodology and the problem they have stated to resolve [Score] *

1. Inappropriate | 3. Appropriate | 5.Very appropriate

*Mark only one oval.*

| 1 | 2 | 3 | 4 | 5 |
|---|---|---|---|---|
| ⬭ | ⬭ | ⬭ | ⬭ | ⬭ |

14. Fit between the project's approach/methodology and the problem they have stated to resolve [Comment, optional]

Please, comment and explain the score you have given. You can also give constructive feedback to help the project improve on this criteria.

15. How relevant is the project to the COVID-19 crisis and alleviation of suffering? [Score] *

1. Not at all | 3. Moderately | 5. Essential

*Mark only one oval.*

| 1 | 2 | 3 | 4 | 5 |
|---|---|---|---|---|
| ⬭ | ⬭ | ⬭ | ⬭ | ⬭ |

16. How relevant is the project to the COVID-19 crisis and alleviation of suffering? [Score]

Please, comment and explain the score you have given. You can also give constructive feedback to help the project improve on this criteria.

17. The project's state of progress [Score] *

1. No results yet | 3. Some results but with a promising development plan | 5. Working proof of concept

*Mark only one oval.*

| | 1 | 2 | 3 | 4 | 5 |
|---|---|---|---|---|---|
| | ○ | ○ | ○ | ○ | ○ |

18. The project's state of progress [Comment, optional]

Please, comment and explain the score you have given. You can also give constructive feedback to help the project improve on this criteria.

19. Clarity and relevance of the project timeline and it's needs for future (major tasks, milestones) [Score] *

1. Unclear and irrelevant | 3. Clear but relevant points are missing | 5. Very clear and relevant

*Mark only one oval.*

| 1 | 2 | 3 | 4 | 5 |
|---|---|---|---|---|
| ○ | ○ | ○ | ○ | ○ |

20. Clarity and relevance of the project timeline and it's needs for future (major tasks, milestones) [Comment, optional]

Please, comment and explain the score you have given. You can also give constructive feedback to help the project improve on this criteria.

21. The project's ability to actively engage and align itself with all the relevant groups and possible stakeholders [Score] *

1. Unaware | 3. Aware but unclear integration | 5. Actively engaged

*Mark only one oval.*

| 1 | 2 | 3 | 4 | 5 |
|---|---|---|---|---|
| ○ | ○ | ○ | ○ | ○ |

22. Ability to actively engage and align itself with all the relevant groups and possible stakeholders [Comment, optional]

Please, comment and explain the score you have given. You can also give constructive feedback to help the project improve on this criteria.

23. How well has the project addressed concerns such as quality testing? [Score] (If applicable)

1. Not well | 3. Clear but not entirely appropriate | 5. Very clear and relevant

*Mark only one oval.*

| 1 | 2 | 3 | 4 | 5 |
|---|---|---|---|---|
| ◯ | ◯ | ◯ | ◯ | ◯ |

24. How well has the project addressed concerns such as quality testing? [Comment, optional]

Please, comment and explain the score you have given. You can also give constructive feedback to help the project improve on this criteria.

25. The scalability of the project in the long term, taking manufacturing into consideration [Score] *

1. No sustainability model and no scaling potential | 3. Sustainable and scalable but still unstructured plan | 5. Already applying a sustainable plan & good scalability

*Mark only one oval.*

| | 1 | 2 | 3 | 4 | 5 |
|---|---|---|---|---|---|
| | ⬭ | ⬭ | ⬭ | ⬭ | ⬭ |

26. The sustainability and scalability of the project in the long term [Comment, optional]

Please, comment and explain the score you have given. You can also give constructive feedback to help the project improve on this criteria.

27. The project's dissemination strategy (quality of documentation for goals, results, methods and needs; open access model, outreach) [Score] *

1. Irreproducible and poorly documented 3. Good documentation but with few weaknesses 5. Easily reproducible & very well communicated

*Mark only one oval.*

| | 1 | 2 | 3 | 4 | 5 |
|---|---|---|---|---|---|
| | ⬭ | ⬭ | ⬭ | ⬭ | ⬭ |

28. The projects dissemination strategy (quality of documentation for goals, results, methods and needs; open access model, outreach) [Comment, optional]

29. Project originality *

How much is app/device this needed? are there already apps or devices that can do this in an open source way? 1. Unoriginal and done before. 3.Done before but not in an open source way. 5. Completely original as far as the reviewer is aware.

*Mark only one oval.*

| 1 | 2 | 3 | 4 | 5 |
|---|---|---|---|---|
| ◯ | ◯ | ◯ | ◯ | ◯ |

30. Originality [Comment, optional]

#### Additional comments and reviewer self-evaluation

You may provide below additional comments addressed to the team leaders of the project, and private comments to the admin team

Please, also assess your expertise in the fields relevant to evaluate the project.

31. [To the team leaders] Other general constructive comments about the project *

32. [To admins] Private comment - anything you would like to inform the admin team of, privately.

33. What is your expertise that is relevant to evaluate this project? *

Please provide justification of your expertise if possible.

34. Would you like us to include you in our "Experts community" and provide similar reviews to projects when you have time in the future? *

*Mark only one oval.*

- Yes
- No
- Maybe

35. What is the reviewers Slack Handle? *

This content is neither created nor endorsed by Google.

Google Forms
